# Clinical and molecular characterization of virus-positive and virus-negative Merkel cell carcinoma

**DOI:** 10.1101/587626

**Authors:** Gabriel J. Starrett, Manisha Thakuria, Tianqi Chen, Christina Marcelus, Jingwei Cheng, Jason Nomburg, Aaron R. Thorner, Michael K. Slevin, Winslow Powers, Robert T. Burns, Caitlin Perry, Adriano Piris, Frank C. Kuo, Guilherme Rabinowits, Anita Giobbie-Hurder, Laura E. MacConaill, James A. DeCaprio

## Abstract

Merkel cell carcinoma (MCC) is a highly aggressive neuroendocrine carcinoma of the skin mediated by the integration of Merkel cell polyomavirus (MCPyV) and expression of viral T antigens or by ultraviolet induced damage to the tumor genome from excessive sunlight exposure. An increasing number of deep sequencing studies of MCC have identified significant differences between the number and types of point mutations, copy number alterations, and structural variants between virus-positive and virus-negative tumors. In this study, we assembled a cohort of 71 MCC patients and performed deep sequencing with OncoPanel, a next-generation sequencing assay targeting over 400 cancer-associated genes. To improve the accuracy and sensitivity for virus detection compared to traditional PCR and IHC methods, we developed a hybrid capture baitset against the entire MCPyV genome. The viral baitset identified integration junctions in the tumor genome and generated assemblies that strongly support a model of a hybrid, virus-host, circular DNA intermediate during integration that promotes focal amplification of host DNA. Using the clear delineation between virus-positive and virus-negative tumors from this method, we identified recurrent somatic alterations common across MCC and alterations specific to each class of tumor, associated with differences in overall survival. Comparing the molecular and clinical data from these patients revealed a surprising association of immunosuppression with virus-negative MCC and significantly shortened overall survival. These results demonstrate the value of high-confidence virus detection for identifying clinically important features in MCC that impact patient outcome.

## Introduction

Merkel cell carcinoma (MCC) is a highly aggressive neuroendocrine carcinoma of the skin. Risk factors for developing MCC include advanced age, light skin color with excessive sunlight exposure, and a variety of immunocompromised conditions (1). In 2008, Merkel cell polyomavirus (MCPyV) was first detected by Southern blot in some but not all MCC tumors with integration of viral DNA occurring at several different chromosomal sites. Importantly, an identical clonal integration pattern was detected in one tumor and corresponding metastatic lymph node (2). This important insight implied that integration of the viral DNA was an early if not initiating event in virus-positive MCC oncogenesis. MCPyV infects most people, typically at an early age, and results in an asymptomatic and lifelong infection indicated by the presence of antibodies to the viral coat protein VP1 (3, 4). Although MCPyV DNA can be readily detected on the skin, the cell types where the virus replicates *in vivo* have not been determined (5).

Since the original discovery of MCPyV, it has become increasingly clear that virus-positive MCC has a different etiology than virus-negative MCC (1). While virus-positive MCC expresses the viral oncogenes Large T antigen (LT) and Small T antigen (ST), the tumor genome usually contains very few mutations in cellular oncogenes and tumor suppressor genes. In contrast, studies using whole exome or targeted hybrid capture sequencing have revealed that virus-negative MCC has an exceptionally high somatic mutation load predominated by UV-mediated mutations with frequent mutations in *RB1*, *TP53*, *NOTCH1*, and *FAT1* (6, 7). Whole genome sequencing (WGS) of MCC confirmed virus-positive MCC exhibits a globally lower, non-UV-mediated, mutation burden as well as few somatic copy number amplifications, deletions, and rearrangements compared to virus-negative MCC, while providing new insights into the structure and mechanism of virus integration (8).

Accurate detection of the presence of MCPyV and distinguishing between virus-positive and virus-negative MCC is important for understanding the oncogenesis and cell-of-origin and potentially for therapeutic options. Currently, there is no routine clinical effort to distinguish between virus-positive MCC and virus-negative MCC. Several recent studies have hinted at differences between virus-positive MCC and virus-negative MCC in presentation, age, and response to immunotherapy (9–15). However, current techniques for determining viral status have yielded either inaccurate or ambiguous results. Although WGS provides much more genetic information on the tumor and viral genome compared to targeted approaches, it remains impractical for clinical evaluation of MCC.

The most common methods for detection of MCPyV in MCC include PCR amplification of MCPyV DNA from DNA isolated from MCC tumors or immunohistochemistry (IHC) staining for MCPyV LT using monoclonal antibodies CM2B4 and Ab3 (16, 17). However, both PCR and IHC have been shown to be unreliable in distinguishing between virus-positive from virus-negative MCC. For example, a recent study of 282 cases of MCC evaluated virus-positivity by IHC with CM2B4 and Ab3 and PCR with a previously validated primer set (18). Notably, there was concordance for all three assays in only 167 of 282 (59.2%) cases with an additional 62 cases positive for 2 of the 3 tests. The remaining 53 (18.8%) were positive for 1 test or none. This study assigned the MCC to be virus-positive if 2 or 3 tests were positive; therefore, detection of viral DNA by PCR alone was not sufficient for a tumor to be called virus-positive MCC. Furthermore, because of the sensitivity of PCR in detecting DNA, a lower limit of 0.01 copy of MCPyV DNA per tumor cell was called virus-positive MCC. Tumors containing < 0.01 viral copies/cell were called virus-negative. A different study using RNA-FISH to detect mRNA specific for MCPyV LT and ST found this method to be as sensitive as qPCR when using two primer sets and the viral copy number was set to ≥ 0.004/cell (19). The AMERCK test detects circulating antibodies against the MCPyV ST (20). The sensitivity of this test is low for detection of virus-positive MCC but, when positive, can be used as a biomarker for disease status (20).

The high somatic mutation burden in virus-negative MCC is predicted to result to yield more tumor neoantigens than melanomas or non-small cell lung cancers (median of 173, 65, and 111 neoantigens/sample, respectively) (21) (22). As observed for other tumor types, the high neoantigen burden in virus-negative MCC corresponds with a higher degree of tumor infiltrating lymphocytes in some tumors, but these tumors also express PD-L1 rendering these lymphocytes ineffective (7). Despite the numerous observed differences in mutation rate and number of predicted neoantigens, both virus-positive MCC and virus-negative MCC tumors have shown high response rates to PD-L1 and PD1 checkpoint blockade therapy (14, 15).

For further advancements to be made in understanding MCC, especially for patients not responsive to current therapies, clear and accurate determination of the MCPyV virus status and actionable variants in these tumors are required. In this study, we developed a viral hybrid capture next-generation sequencing (NGS) method to detect the presence of integrated MCPyV DNA in FFPE clinical specimens. This approach was combined with targeted sequencing of several hundred cancer-related genes to assess oncogenic changes in the tumor genome. We compared this viral hybrid capture approach to PCR detection of viral DNA, IHC for detection of MCPyV LT, and synoptic assessment of MCC pathology.

## Results

### Summary of patient cohort

A total of 71 patients diagnosed with MCC were included in this study (Table 1). The median (95% CI) follow-up duration from initial diagnosis of MCC was 47 (95% CI: 38 - 60) months based on inverse Kaplan-Meier estimation. Overall, 69 enrolled patients were white and two were black. Forty (56%) patients were male. The median age was 70 years (range: <50 to 93). The initial site of MCC presentation was in the head and neck (27%), upper extremity (20%), lower extremity (21%) and trunk (32%). The seventh edition TNM staging system of the American Joint Committee on Cancer (AJCC) was used to classify the initial presentation of MCC with 27% presenting at stage I, 14% Stage II, 42% Stage III, and 17% Stage IV.

**Table 1.** Patient Characteristics (N=71)

| **Characteristics** | **All (N=71)** |
| --- | --- |
| **Age at initial diagnosis, yrs** | 70 (10-93) |
| **Age at initial diagnosis, yrs** |  |
| ≤70 | 36 (51%) |
| >70 | 35 (49%) |
| **Gender** |  |
| Female | 31 (44%) |
| Male | 40 (56%) |
| **Race** |  |
| Black or African American | 2 (3%) |
| White | 69 (97%) |
| **Initial site** |  |
| Head | 19 (27%) |
| LE | 15 (21%) |
| Trunk | 23 (32%) |
| UE | 14 (20%) |
| **AJCC stage at initial diagnosis** |  |
| I | 19 (27%) |
| II | 10 (14%) |
| III | 30 (42%) |
| IV | 12 (17%) |
| **Significant immunosuppression** |  |
| No | 61 (86%) |
| Yes | 10 (14%) |
| **Prior chemotherapy or radiation** |  |
| No | 53 (75%) |
| Yes | 18 (25%) |

### Somatic Variant Analysis of Targeted Sequencing

All 71 patients underwent OncoPanel analysis (23). Genomic studies were performed using DNA isolated from tumors obtained at the time of initial diagnosis (50) or upon relapse (21). The total number of mutations ranged from 0 to 73 corresponding to a Tumor Mutational Burden (TMB; mutations/megabase) from 0 to 38.89 with four cases containing no detectable mutations (Figure 1A, Table S1). From this mutation data, patients were binned into TMB-high (>=20), TMB-intermediate (>6<20), and TMB-low (<=6). A limited set of mutation signatures could be identified (see Methods). The UV mutational signature (Signature 7) was detected in 24 cases, corresponding to the TMB-high patients (24). Additional mutational signatures identified included Aging (Signature 1; 3 cases), APOBEC (Signatures 2 & 13; 4 cases with 3 that also had an UV signature), and Signature 5 (one case) (Figure 1A, Table S1). TMB had some correlation with the number of copy number altered genes (Figure 1B). Several genes including *RB1*, *TP53*, *KMT2D*, *NOTCH1*, *NOTCH2,* and *FAT1* were highly enriched for missense and truncating mutations (Figure 1C, Figure S1). Single and dinucleotide substitutions in *RB1* and *TP53* revealed that most were likely mediated by UV damage (CC>TT, C>T; Figure 1D).

**Figure 1.**
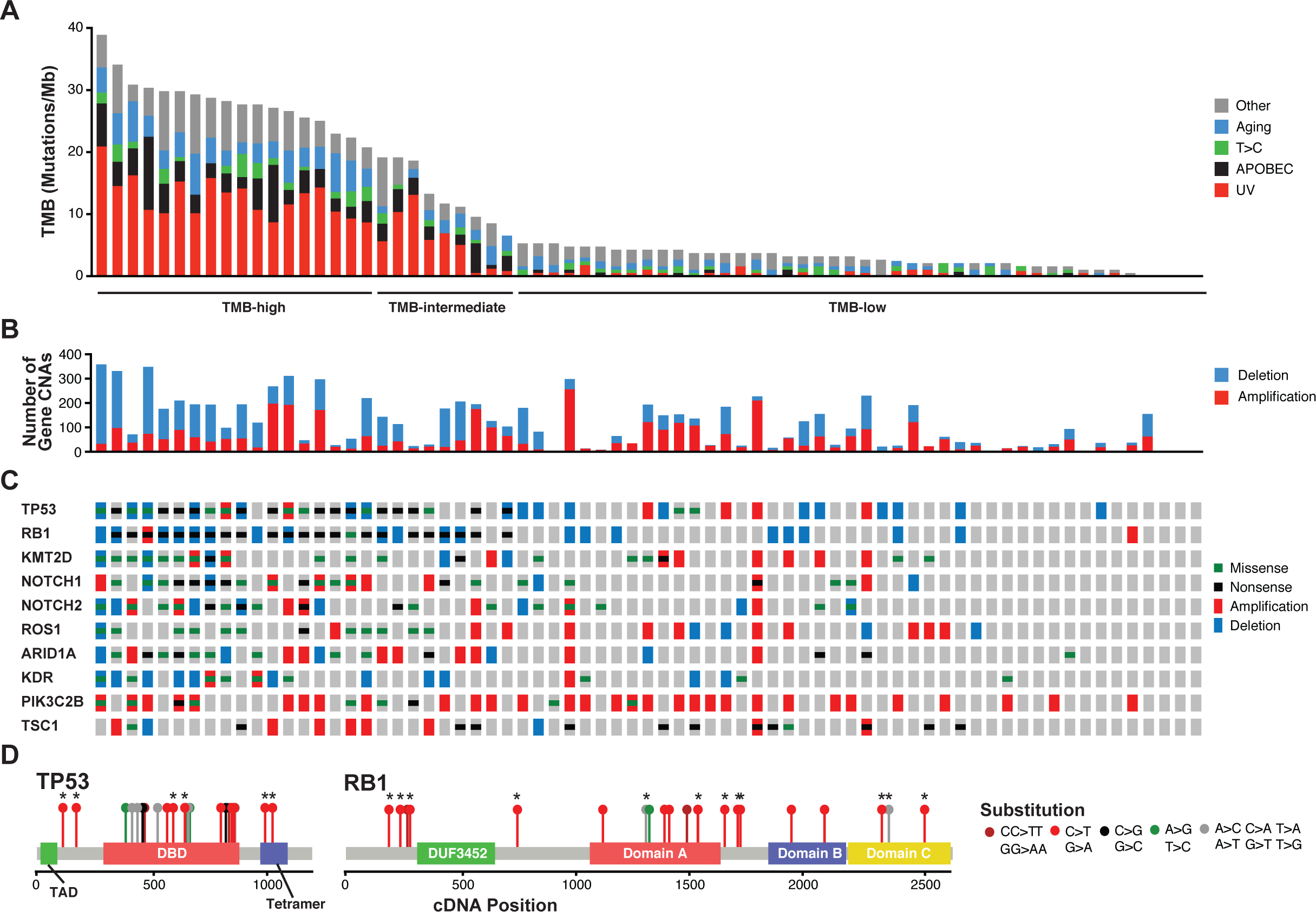
Somatic variants in Merkel cell carcinoma. A. Tumor mutation burden (TMB) for each patient in descending order colored by mutation signature. B. Count of gene copy number alterations per patient. C. OncoPrint for the top 10 genes with the greatest number of point mutations in this MCC cohort. D. Distribution of point mutations in the cDNA of RB1 and TP53 from this MCC cohort. Functional domains of p53 and pRB are highlighted by colored boxes. Each type of base substitution is highlight by a different color lollipop and nonsense mutations are indicated by asterisks.

Copy number variants (CNVs) were examined individually as well against each other and other likely functional somatic changes for significant co-occurrence or mutual exclusivity (Table S2). Clusters of significantly co-occurrent CNVs were determined via network analysis (Figure 2A, Table S3). From these analyses, two chromosomal regions were found to be altered in more than 36% of cases (Figure 2B & C). Chromosome 10 (cluster 14) had frequent copy number loss with many tumors showing complete loss of the entire chromosome (Figure 2B) (25). Some cancer-relevant genes on chromosome 10 include PTEN and SUFU, negative regulators of PI3K and Hedgehog signaling respectively, with deletions reported in prior studies of MCC (25, 26). A region of Chr1q (cluster 13) was amplified in 28 cases. This region includes MDM4 (also known as MDMX), whose protein product cooperates with MDM2 to promote the ubiquitination and subsequent degradation of p53 (Figure 2B) (27, 28). In addition, we observed a focal amplification of MYCL in Chr1p (cluster 4), which was reported in an earlier study of MCC (29).

**Figure 2.**
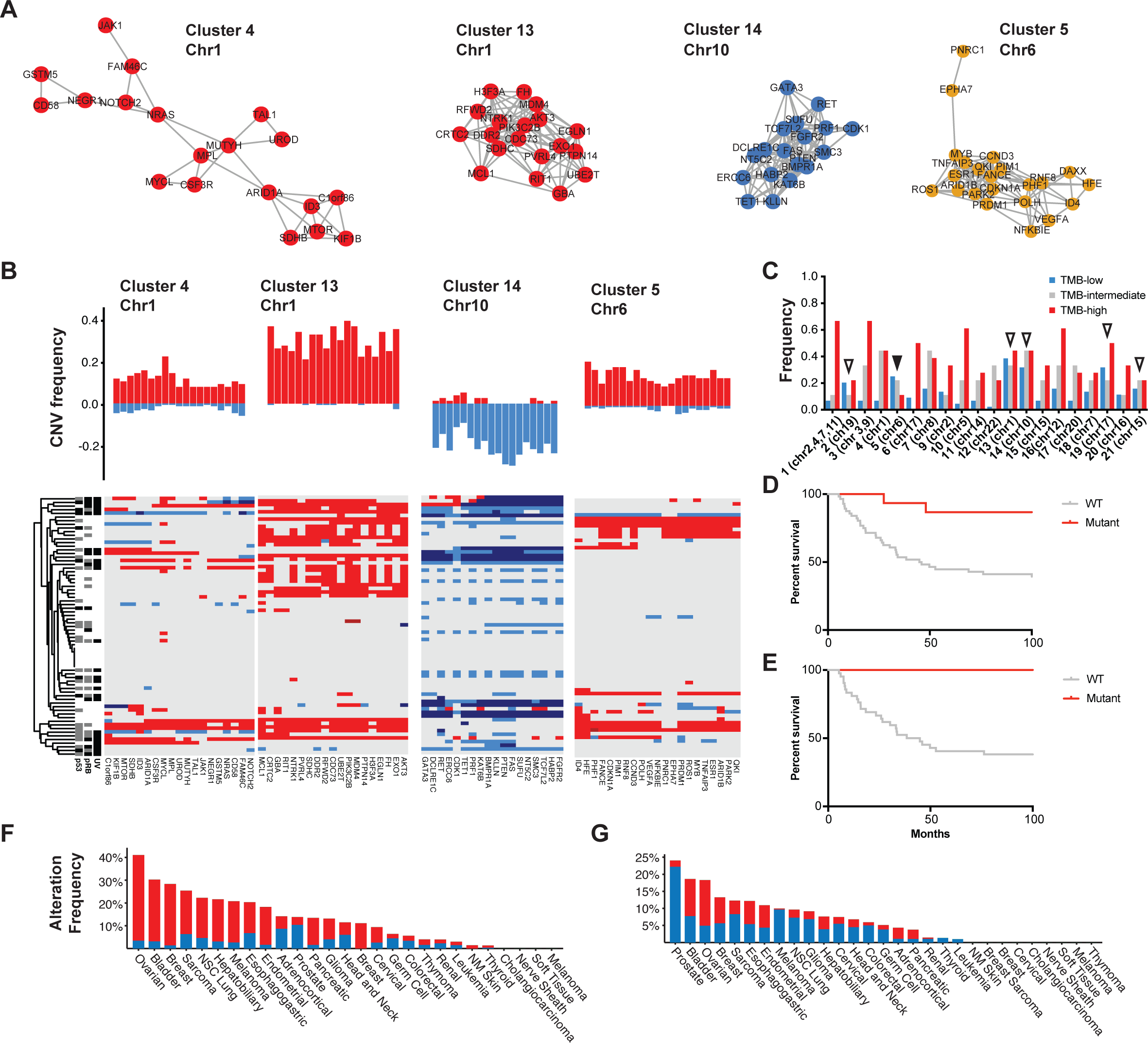
Recurrent copy number variants in MCC. A. Representative network analysis clusters of significantly co-modified genes in MCC on chromosomes 1 (red), 6 (yellow) and 10 (blue). B. Frequency of amplifications (red) and deletions (blue) for the genes comprising representative CNV clusters and their occurrence in each patient with UV, RB1, and TP53 status clustered by all variants. C. Counts of each CNV cluster colored by TMB-low (blue), - intermediate (grey), and -high (red) categories. Clusters that are nearly equivalent between TMB-low and -high (<2:1 ratio are highlighted by open triangles). The cluster that is more frequent in TMB-low than -high is highlighted by a black-filled triangle. D. Kaplan-Meier plot of overall survival stratified by chromosome 6 amplification for all patients. E. Kaplan-Meier plot of overall survival stratified by chromosome 6 amplification for primary tumors. F-G. Analysis of TCGA cancers for the two most abundant CNV clusters (13 & 14, respectively) in MCC.

The CNV clusters were observed at nearly equal frequencies in both TMB-high and TMB-low cases (Figure 2B & C). Many of the other CNV clusters were strongly associated with UV signature and high TMB (Figure 2C). However, cluster 5 that included an amplification of chromosome 6 was more than twice as frequent in TMB-low tumors than TMB-high tumors (Figure 2C). Interestingly, 35% of metastatic tumors carried cluster 5 and all but one of these metastatic tumors were TMB-low MCC. Furthermore, CNV cluster 5 was 2.5 times more frequent in TMB-low (22.2%) than TMB-high (8.7%) tumors in primary tumors. Both TMB-low and TMB-high patients with amplification of CNV cluster 5 had significantly improved overall survival compared to wild-type carrying patients that had a median survival time of 41.65 months (p=0.0027). Restricting this analysis to only primary tumors, revealed that there were no deaths at the time of this study in patients carrying this amplification (p<0.001) (Figure 2D & E). The recurrent copy number events on chromosomes 1 and 10 were compared within TCGA for similarities to other tumor types (Figure 2F-G). This analysis revealed that the chromosome 1 amplification was also frequently observed in ovarian, breast, and bladder cancers; whereas, the chromosome 10 loss was most frequently seen in prostate cancer.

### Analysis of Viral Sequences in Tumors

Of the 71 tumors analyzed by Oncopanel, 48 were re-analyzed by Oncopanel (Profile/OncoPanel version 3, POPv3) combined with a hybrid-capture probe bait set targeting the entire genome of MCPyV (VB2). For the 48 cases, the number of MCPyV reads ranged from 0-21,095,751 with only a single case having zero MCPyV reads (Figure 3A). In total, 28 cases had substantial reads (>6,800) mapping to the MCPyV genome that also supported integration of the virus into the host genome. For the remaining 20 cases without evidence for integration, the number of viral reads ranged from 0 to 971. Generally, these cases had reads that covered less than 10% of the viral genome with the normalized coverage less than two logs compared to samples with evidence for virus integration (Figure 3B & C, Table S3). Concordantly, the viral reads from most of these cases were unable to be assembled into larger viral contigs. Two cases, MCC011 and MCC015, had 212 and 177 MCPyV reads that could be assembled into nine and five contigs each smaller than 761 base pairs, respectively. Case MCC007 had the most reads of any likely virus-negative sample and could be assembled into a single 5343 bp contig. However, analysis of the point and deletion variants in these aforementioned viral contigs revealed that they were identical to the virus sequence from patient MCC037 indicating that the viral reads resulted from low level contamination (<0.005% of MCC037 MCPyV reads were detected in other samples).

**Figure 3.**
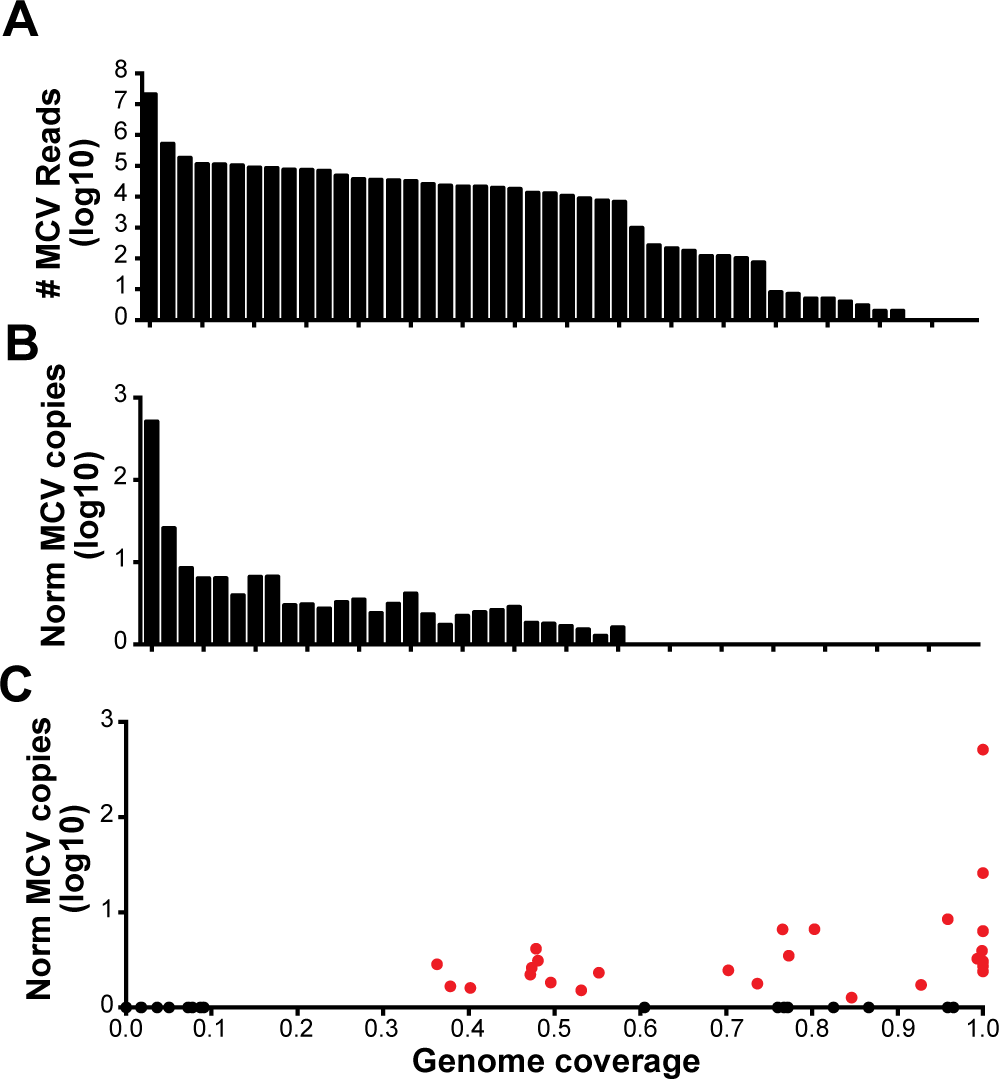
Detection of MCPyV via targeted capture and NGS. A. Raw number of reads mapping to the MCPyV genome per patient from VB2 (n=48). B. Normalized count of MCPyV reads based on number of human reads and fraction of viral genome covered. C. Scatter plot of genome coverage vs normalized MCPyV copies with virus-positive patients highlighted in red and virus-negative patients in black.

For the 28 cases with evidence for integration of the viral DNA into the tumor, the number of reads mapping to the viral genome ranged from 6,824-21,095,751 (Median: 28,726). Consistent with previous reports, the integrated viral genome had undergone extensive mutagenesis with large deletions (>100 bp) particularly in the 3’ half of LT (Figure 4). In 10 cases, approximately half of the total viral genome was deleted, 6 cases had approximately 25% of the viral genome deleted, while 12 cases had sequences corresponding to the entire or nearly complete genome (Figure 3C & 4). In all but one of the cases with a nearly complete coverage of the viral genome, there was a clonal point mutation which inserted a premature stop codon in LT resulting in truncated proteins between 208 and 771 amino acids (Figure 5A). In a single case (MCC054), LT was truncated by a 5bp deletion resulting in a frame shift putting a premature stop codon in frame. In all cases, the non-coding control region, the N-terminal 208 residues of LT, and an intact ST region of the viral genome were conserved. Beyond indels and nonsense mutations, LT also carried numerous clonal missense mutations (Figure 5A). In stark contrast, ST only had missense mutations at three residues, and the amino acid change A20S is consistent with a previously observed MCPyV strain difference (GenBank identical protein accession number: ACI25295.1). The other missense mutations occurred clonally at H41Y and N100S once in the entire cohort (Figure 5B). Neither of these mutations are present in any of the ST sequences in GenBank.

**Figure 4.**
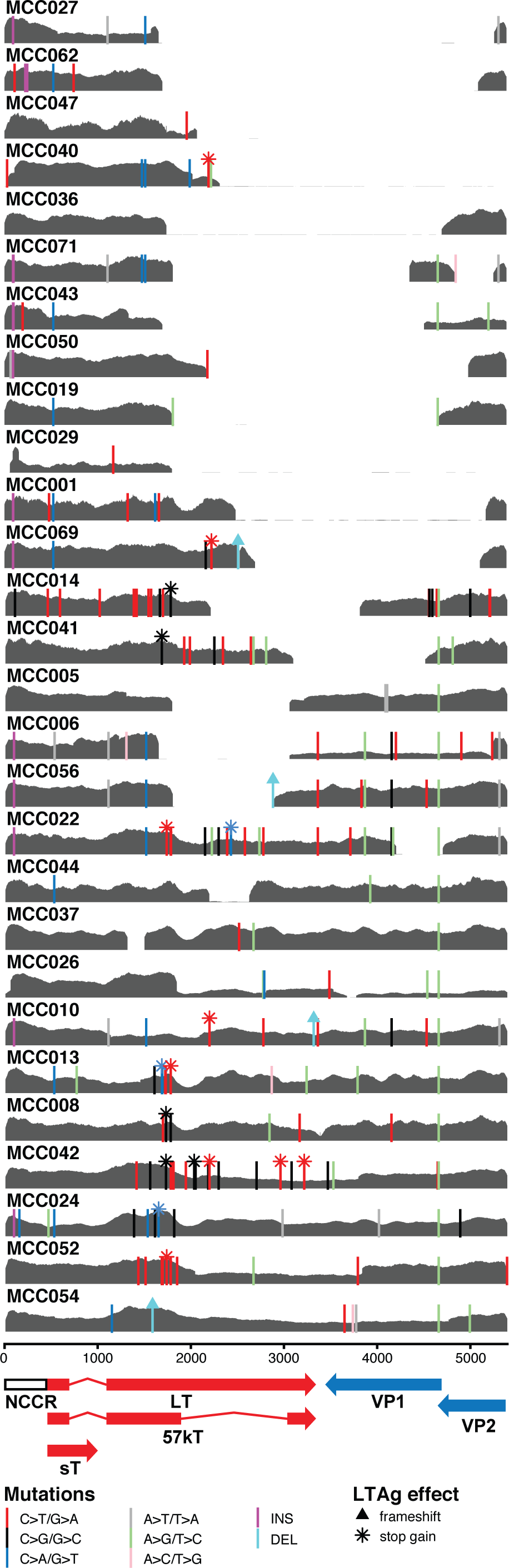
MCPyV coverage and mutations from virus-positive cases. Read coverage for MCPyV in gray and each plot represents a single patient with their ID in the upper left corner. Scales for the coverage plots are set from 0 to the maximum read coverage per patient. Point and insertion-deletion mutations are indicated by vertical lines located at the start point of the mutation colored by the type of base substitution. The effects of point mutations within LT antigen are indicated by a triangle (frameshift) or asterisk (stop gain) at the top of the vertical line of the mutation.

**Figure 5.**
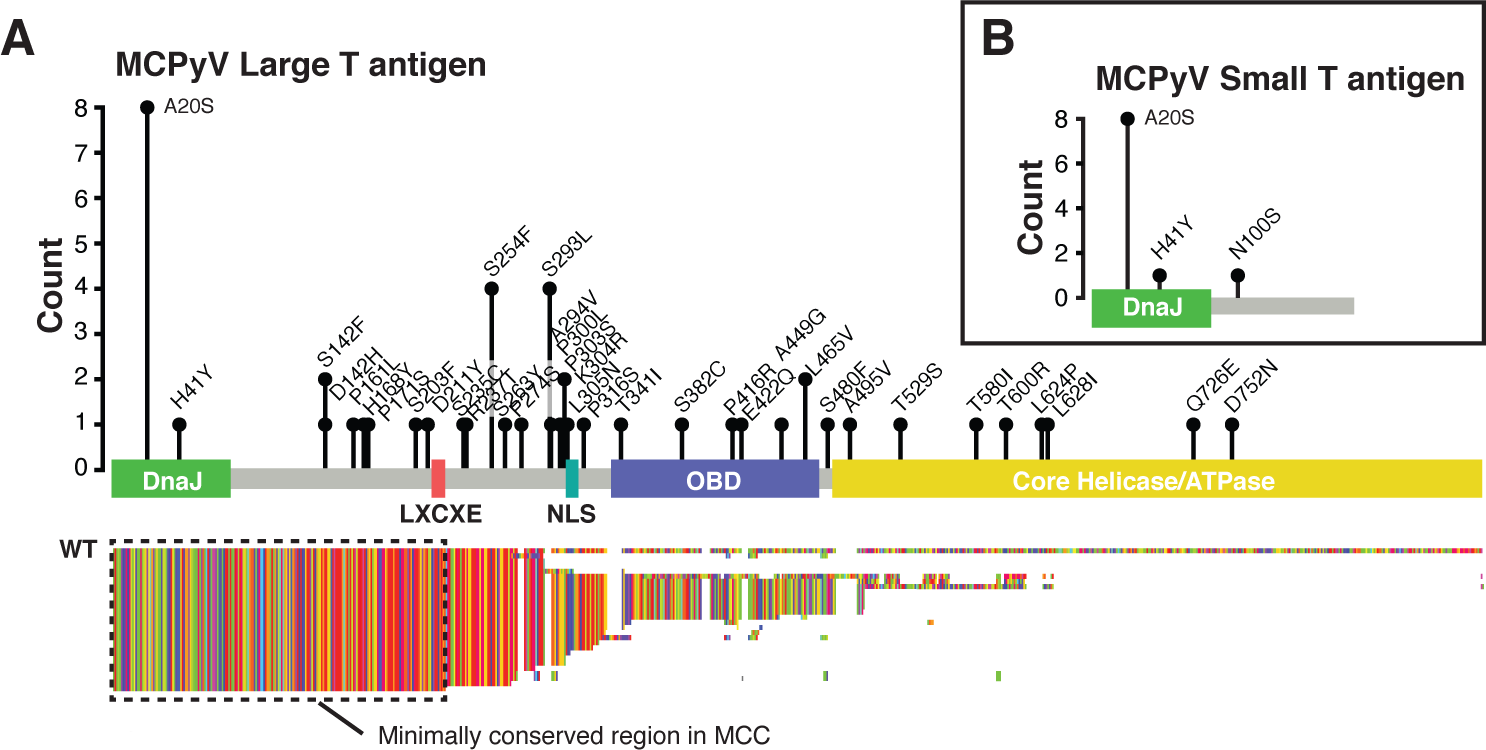
Protein changes in Large and Small T antigens in MCC. A. Lollipop plot of all LT missense mutations relative to the NC_010227.2 MCPyV reference with height reflecting the number of observations in our cohort and residue change labeled above the position. LT domains are highlighted by colored boxes. Below the LT diagram, MAFFT alignment of predicted LT sequences from all virus-positive cases colored by amino acids. B. Lollipop plot of all ST missense mutations relative to the NC_010227.2 MCPyV reference genome

The integration sites were mapped using the oncovirus tools suite (https://github.com/gstarrett/oncovirus_tools) (Figure 6A, Table 2). As previously reported, integrations primarily fell into two categories: either those that appear as a single integration event or as two events separated by >10 kilobases (kb) (8). Interestingly, two cases had integration events in non-identical but overlapping sites in chromosome 1 (Figure 6B). These represent the first reported cases of recurrent viral integration sites or hotspots in MCC.

**Figure 6.**
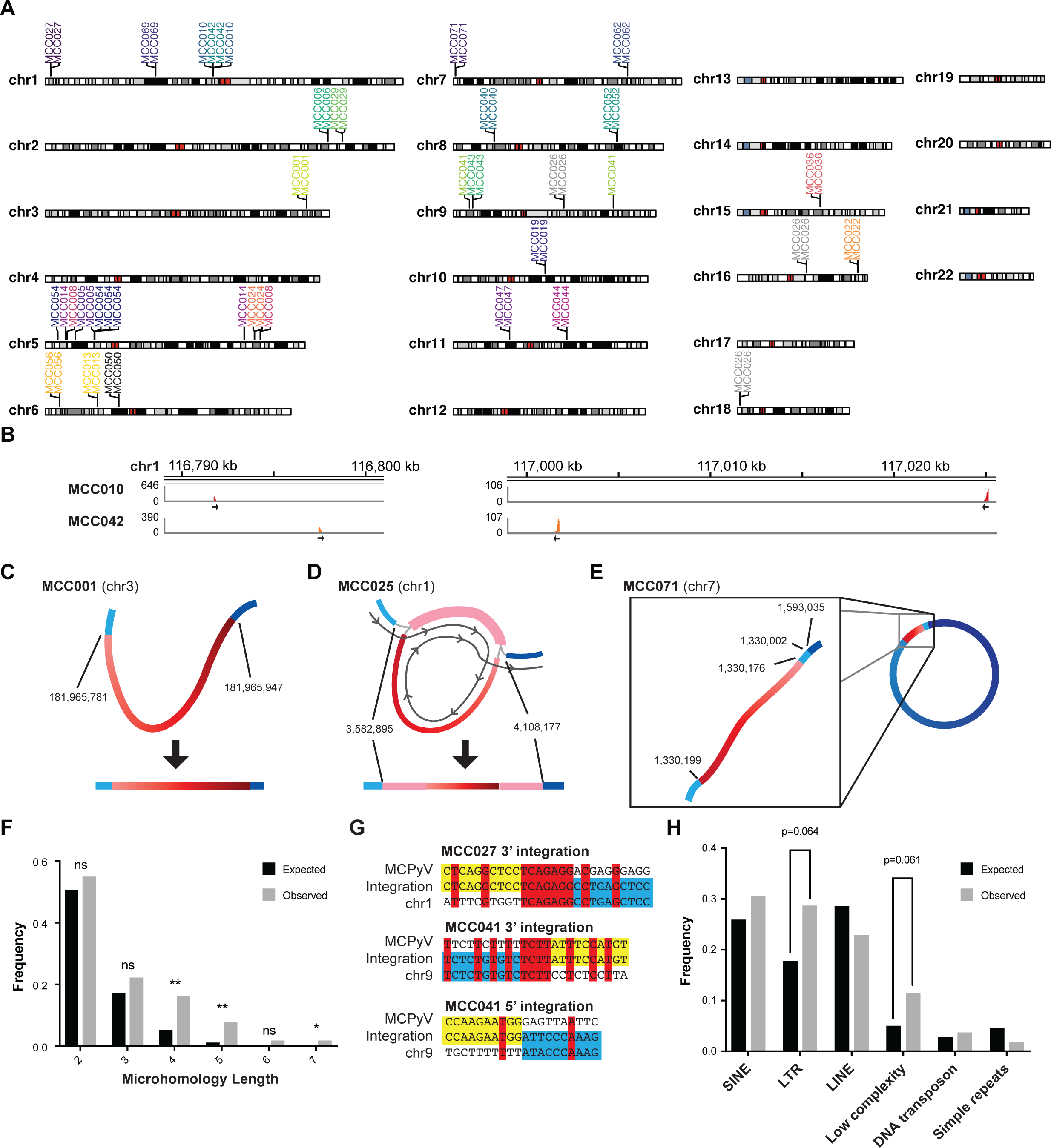
Characterization of MCPyV integration sites. A. Location of integration events in the human genome labeled and colored by patient. B. Coverage of reads corresponding to predicted overlapping integration sites on chromosome 1. Direction of virus-to-host fusion is shown by black arrows. C-E. Representative assembly graphs for different types of viral integrations. Human DNA is a blue gradient and viral DNA is a red gradient representing different genomic segments. Human chromosome positions at the virus junctions are shown. Detailed assembly graphs for all virus-positive cases are in Figure S4. C. Representative single linear assembly graph for integrated MCPyV from case MCC001 on chromosome 3. D. Representative assembly graph of partially duplicated MCPyV genome integrated into the tumor genome of MCC025 on chromosome 1. Path for linearization of assembly graph shown by the dark grey line. E. Representative assembly graph of MCPyV genome integrated into chromosome 7 of MCC071 supporting a circular DNA intermediate diagrammed on the right. F. Barplot showing the frequency of microhomology lengths between two and seven basepairs. Expected values are in black and observed are in grey. Asterisks representing p-values from Fisher’s exact test are represented above the bars (* <0.05, ** <0.01). G. Diagram of representative integration sites with viral sequence highlighted in yellow and host sequence in blue. Matching bases between host and virus are in red. H. Barplot showing the frequency of repetitive elements within 2 kb of integration sites. Expected values are in black and observed are in grey. P-values from Fisher’s exact test are represented above the bars.

**Table 2.**
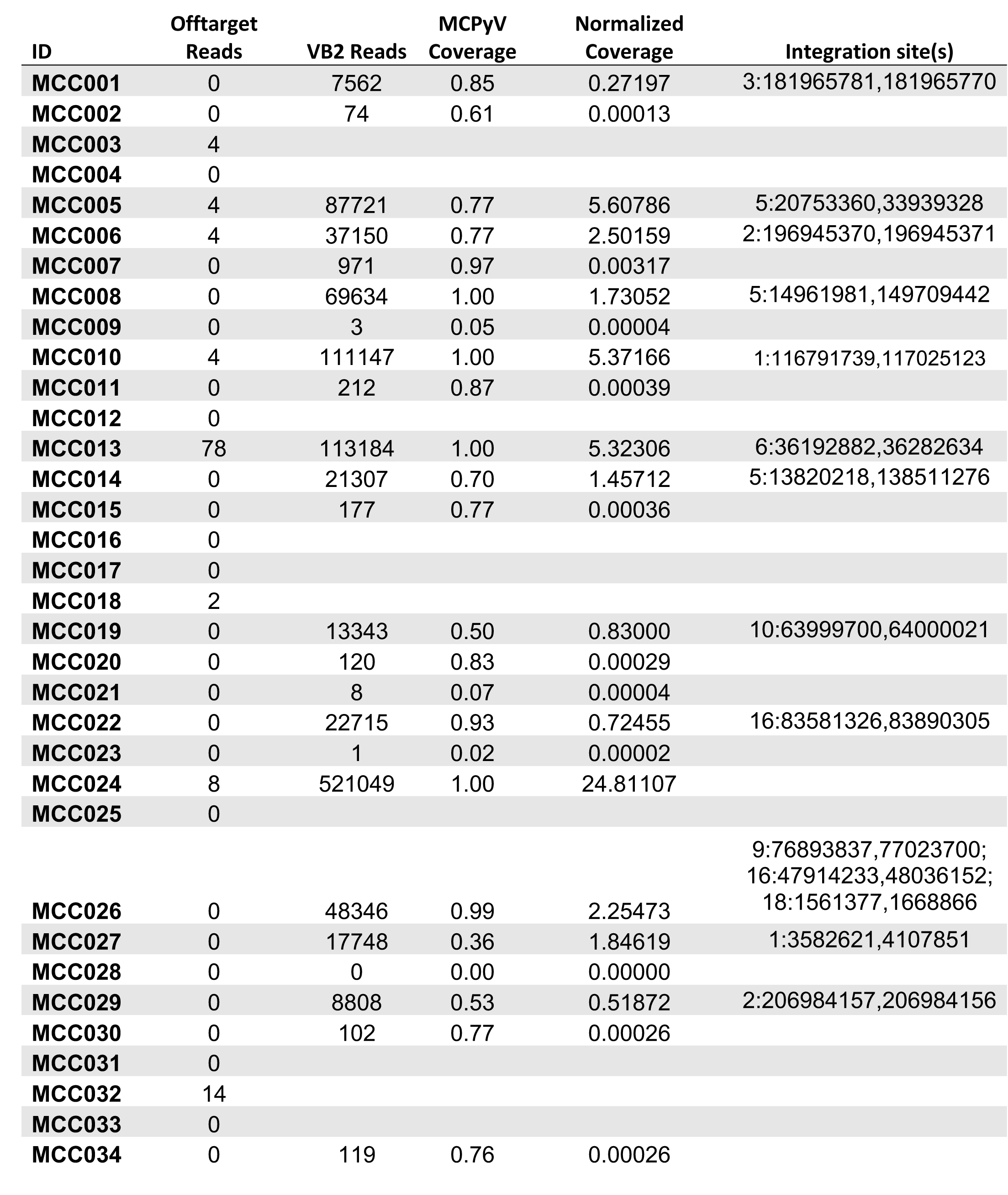
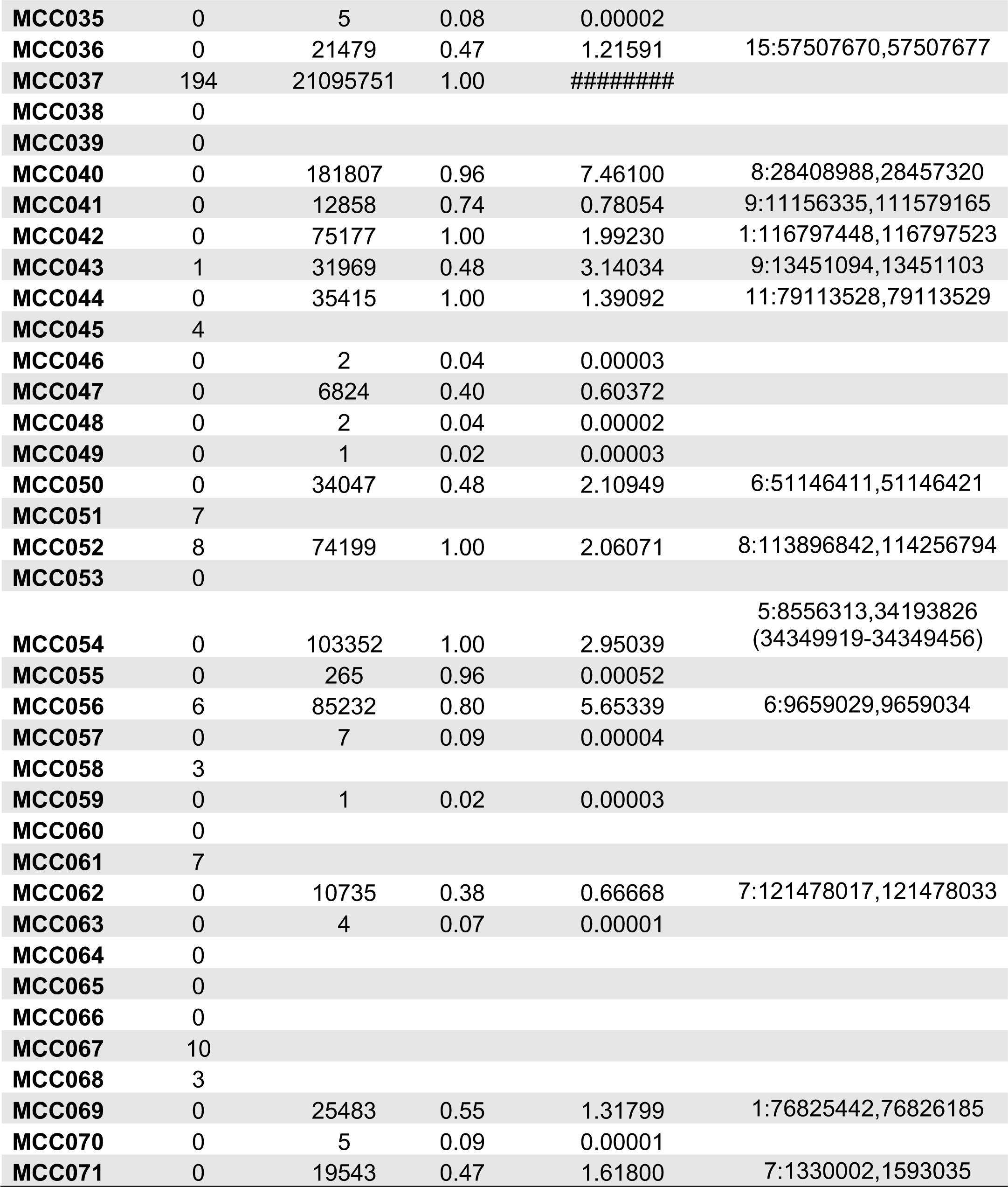
MCPyV Integration sites.

Because of the limited number of capture targets of the OncoPanel platform, determining the extent of virus-mediated amplifications expected in the tumor genome was not possible. However, using the normalized viral coverage, the estimated number of viral genome copies ranged between 1 and 1881 copies (median: 7, interquartile range (IQR): 4-13) (Table 2). Additionally, based on previous observations, we can infer that the regions between viral integration sites were amplified (8). When annotating these regions, we observed that they frequently contain enhancer regions that may contribute to oncogenesis as seen in HPV-associated tumors (30). Patient MCC026 had integrations on chromosomes 9, 16, and 18, all of which had integration sites separated by 107.5 and 129.9 kbp. Using automated computational methods, we could not confidently determine an integration site for case MCC037 with the highest viral genome copy number in this study. Manually interrogating the human sequence hits from the assembly revealed that it matched a tandem repeat sequence flanked by MLT1H2 ERVL-MaLR elements. Based on the estimated copy number and the assembly graph, the viral component of this fusion DNA structure is likely larger than 10 Mbp (Figure S4).

With the high depth of coverage facilitated by the targeted NGS method, high resolution assemblies for the integrated virus were generated. Many integrations that appeared as a single linear contig contained a single copy of the viral genome flanked by the host genome (Figure 6C Figure S4). However, other integrations generated more complex assembly graphs with a multiple contigs linked together in a “pigtails” conformation (Figure 6D, Figure S4). Based on coverage and conformation, this graph likely represents an integration event containing partially duplicated viral genomes fused to different segments of the human genome. For samples with distant integration sties, the directionality of the virus-host junctions strongly supports a circular virus-host DNA fusion intermediate prior to reintegration into the host chromosome. This model is further supported by assemblies in which one arm of the fusion contains sequences from both distant sites of the human genome (Figure 6E, Figure S4).

To address a possible mechanism for integration, we looked for microhomology between the human and MCPyV genomes at fusion junctions. We found significant enrichment for 4, 5, and 7 bp sequence microhomology at the site of integration compared to randomly selected sites in the human and MCPyV genomes (Figure 6F). There was no significant increase in overall homology between MCPyV and human DNA at integration sites versus randomly selected sites. Patient MCC027 had the integration site with the longest stretch of homology and MCC041 had both the integration site with the greatest overall homology on its 3’ end and lowest homology with no microhomology greater than 1bp on its 5’ end (Figure 6G). Additionally, we annotated integration sites for proximity to repeat elements, including LINES, SINES, LTR retrotransposons, and simple repeats in the human genome. No type of repeat element was significantly enriched, but all integration sites were within 1.5kb of a repeat element and there was a trend towards integrations near LTR retrotransposons and low-complexity regions (Figure 6H).

### Distinguishing virus-positive MCC from virus-negative MCC using somatic variants in comparison to Immunohistochemistry and PCR

Given the striking differences in the number of mutations and mutational signature we observed in the VB2 dataset that strongly correlated with virus integration, we compared the data from the OncoPanel and POPv3VB2 datasets to determine the viral status of all 71 tumors studied (Table 3). From the OncoPanel sequencing, we identified off-target reads for MCPyV in a total of 18/71 cases, ranging from 1 to 194 reads total. When compared to the VB2 data, there was a rough correlation between the number of off-target reads and the number of MCPyV reads in the VB2 dataset. There were 8 samples with MCPyV reads in the OncoPanel dataset that were not also analyzed by VB2. None of these 8 cases have any evidence for a UV mutational signature.

**Table 3.**
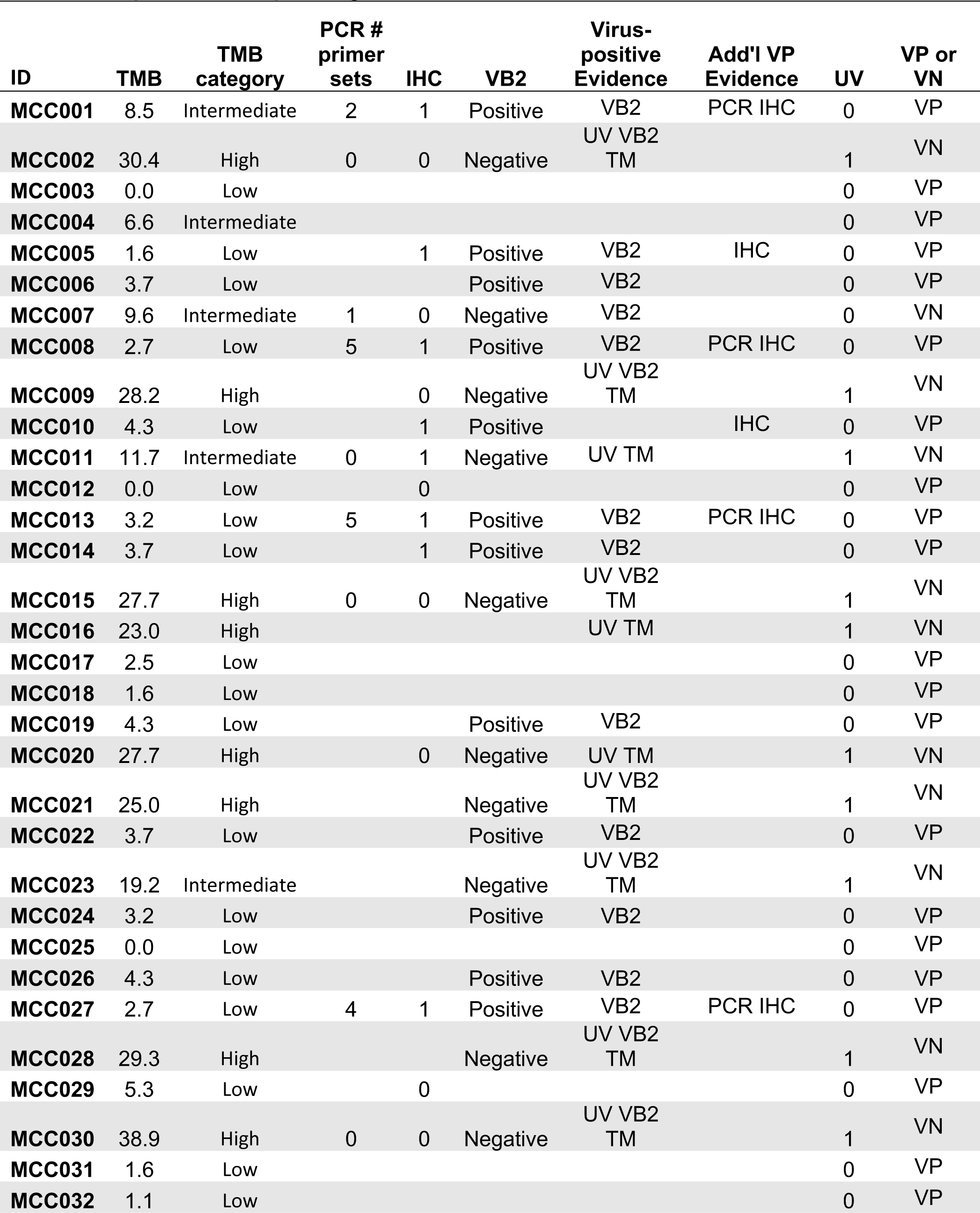
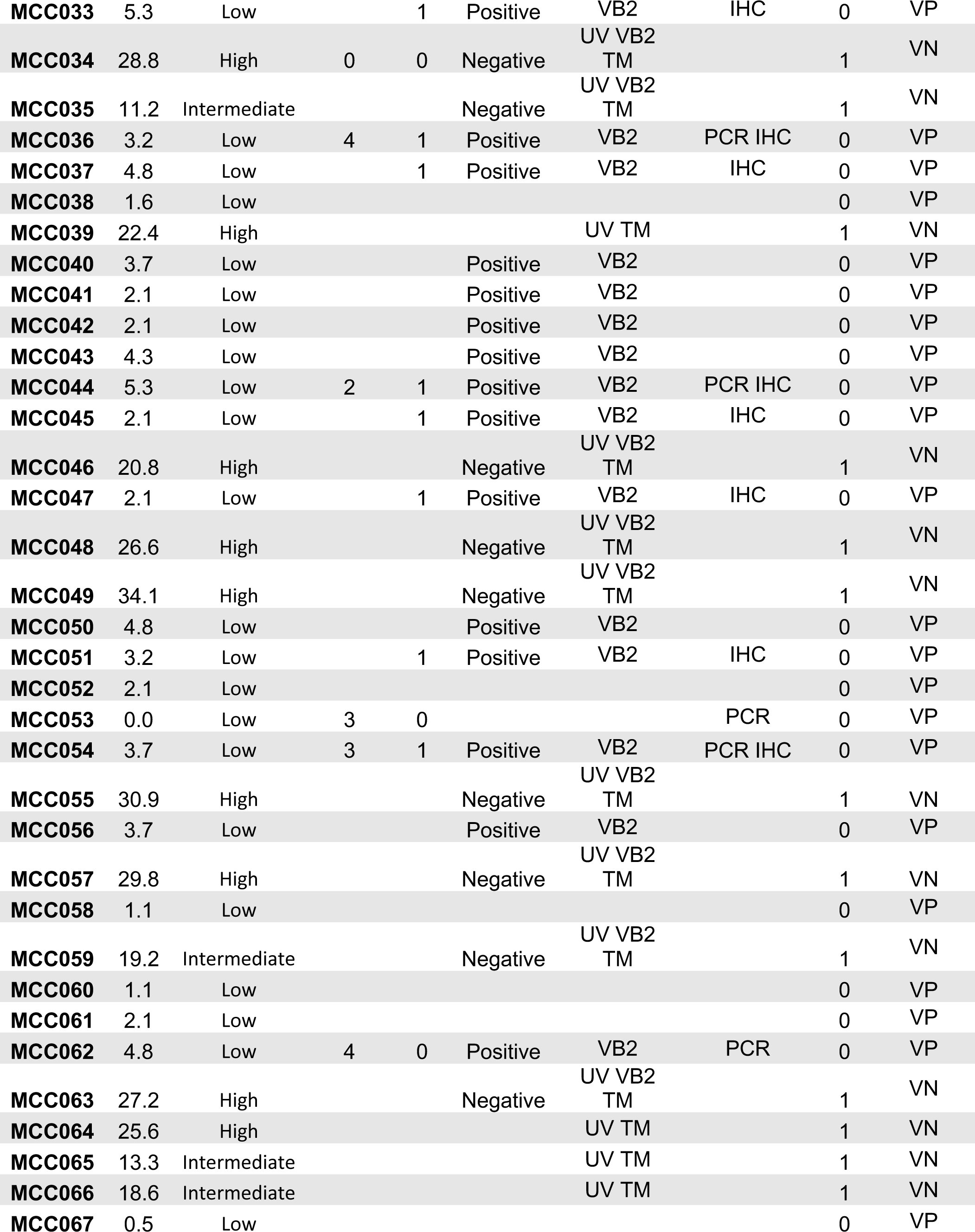
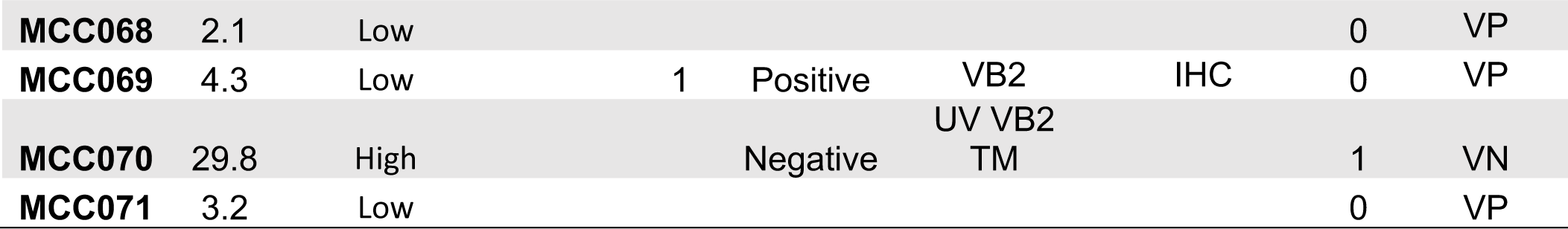
Comparison of sequencing, PCR, and IHC for determination of tumor viral status.

We assessed the total number of mutations, TMB, UV signature, and detection of MCPyV reads to characterize each tumor as virus-positive MCC or virus-negative MCC. Using these criteria, we called 25 tumors as virus-negative. All but one of the virus-negative MCC tumors had a UV mutational signature, had higher number of total mutations (18–73), higher TMB, and absence of integrated MCPyV compared to virus-positive MCC. The virus-negative MCC without a UV signature (MCC007) originally presented as a subcutaneous breast mass (31). A total of 46 MCC tumors of the 71 analyzed were characterized as virus-positive. These virus-positive MCC had an absence of UV mutational signature, a lower number of total mutations (0–16), and lower TMB than did any of the virus-negative MCC. The TMB-low and -high categories had perfect concordance with virus-positive and -negative MCC determined by sequencing, respectively. The TMB-intermediate samples were mostly virus-negative (7/9), but the lowest two TMB patients in this category are likely virus-positive based on VB2 sequencing and absence of UV mutation signature.

FFPE sections were available for 28 of the 71 cases to assess for MCPyV LT by IHC with antibodies CM2B4 and Ab3. For 8 of the virus-negative MCC, all were negative by IHC with both antibodies. For 20 virus-positive MCC cases, we observed 16 stained positive with both antibodies and 4 were negative (Table 3). In addition, DNA was tested by PCR with 5 primer sets for 15 cases. In 9 virus-positive MCC cases, all returned positive results with 2 to 5 primer sets (Table 3). For 6 virus-negative cases, PCR was negative for 5 primer sets and one was positive with one primer set. Interestingly, the virus-negative MCC (MCC007) with one PCR primer set positive also ranked at the TMB borderline (9.58) between virus-negative and virus-positive and did not score as having a UV mutational signature; rather, the majority of mutations were classified as APOBEC-associated.

A synoptic review of dermatopathology was available for 19 cases (Table S4) (32). Criteria evaluated included procedure, laterality, site, size (mm), thickness (mm), lymphovascular invasion, tumor extension, mitotic rate, tumor infiltrating lymphocytes (TILS), growth pattern, neurotropism, and necrosis (%). TILS were largely absent in both virus-positive and virus-negative samples. An infiltrative growth pattern was observed in virus-positive MCC and nodular or nodular infiltrative observed in both forms of MCC. Neurotropism was present in three cases of virus-positive MCC and necrosis ranged from 0 to 40%.

### Statistical comparison of clinical and molecular characteristics

Overall, 28 patients remained disease free after initial therapy and 43 developed one or more relapses or persisted as stage IV. According to the biopsy type and first recurrence status, patients could be grouped into primary biopsy with no further recurrence (N=30), primary biopsy with further recurrence (N=22), and recurrence biopsy (N=19). For all samples annotated with recurrence, the first recurrence occurred before the biopsy was obtained. Regardless of the biopsy type, patients were grouped into no relapse (N=30) and relapse (N=41). **Table 4** shows the association between relapse and genomic characteristics. Among 71 patients, 30 (42.3%) patients had no relapse and 41 (57.7%) had relapse after initial diagnosis. From the Fisher’s exact test results, UV, RB1 status, TP53 status, and virus status were all not significantly associated with relapse (Table 4). If the OncoPanel data obtained after relapse (and prior treatment) was excluded and restricted to the 52 patients with primary biopsy, UV, RB1 status, TP53 status, and virus status were all not significantly associated with relapse (Table S5).

**Table 4.**
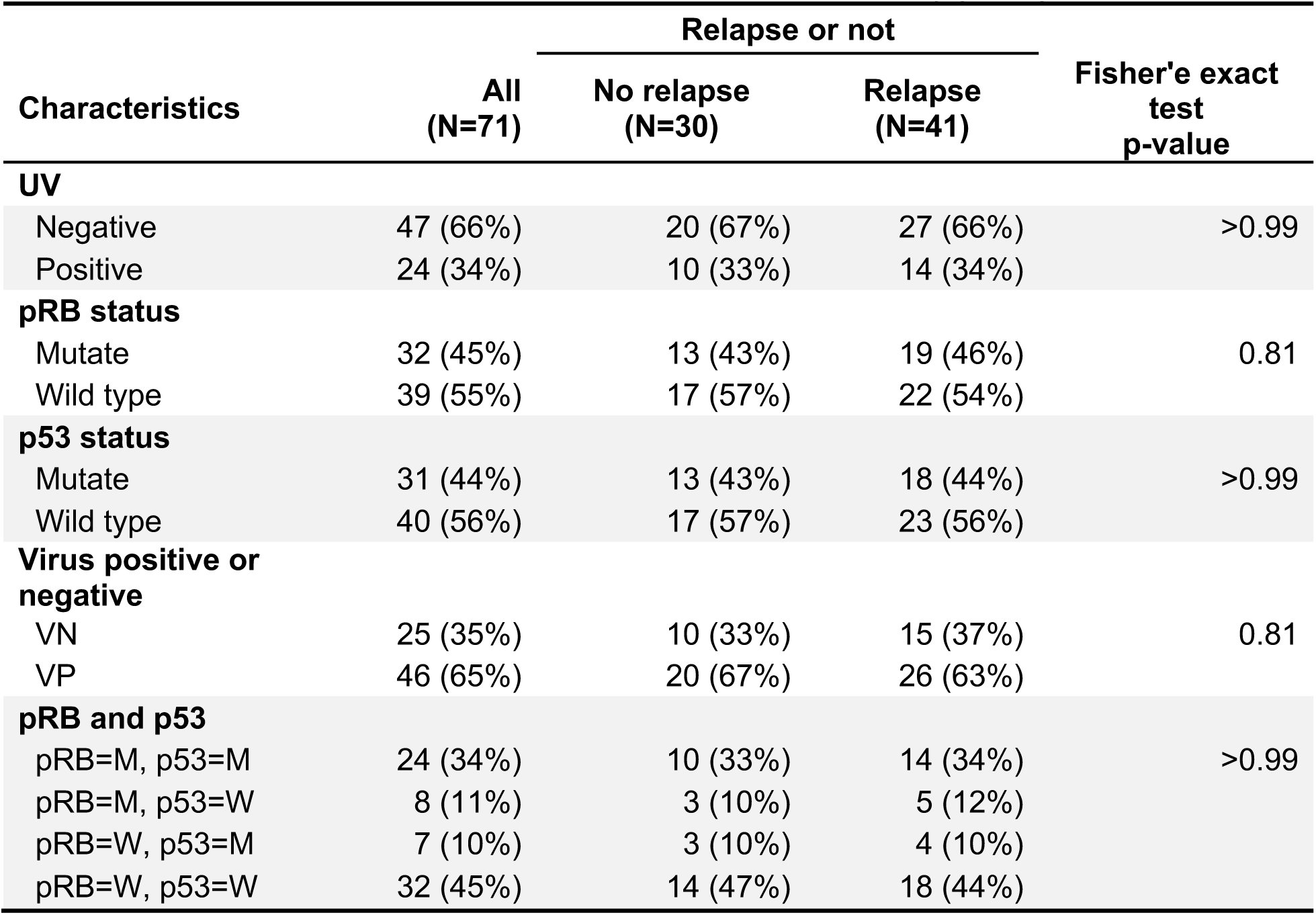
Association between Relapse and Genomic Sequencing (N=71)

Consistent with known risk factors of MCC, 10 of the 71 cases had immunosuppression diagnosed prior to developing MCC. Remarkably, 8 of the 10 (80%) of the immunosuppressed cases were identified as virus-negative MCC with relatively high TMB compared to the 28% virus-negative MCC in immunocompetent patients (Figure 7A, Table 5). Virus-negative MCC was present in three patients with solid organ transplantation, three with auto-immune disease including myasthenia gravis, rheumatoid arthritis, or granulomatosis with polyangiitis, one with monoclonal gammopathy of undetermined significance (MGUS), and another with Waldenstrom’s macroglubulinemia. In contrast, virus-positive MCC was identified in a patient with mantle zone lymphoma having been treated with Rituximab for 3 years and another with somatic mutations in NF1 and GATA2 (33). The median OS for patients with immunosuppression was 17.5 months (95% CI: 5.6-24.4) months, significantly shorter than patients without immunosuppression (48.5 months, 95% CI: 35.4-113.3, p<0.01) (Figure 7B, Table 5).

**Figure 7.**
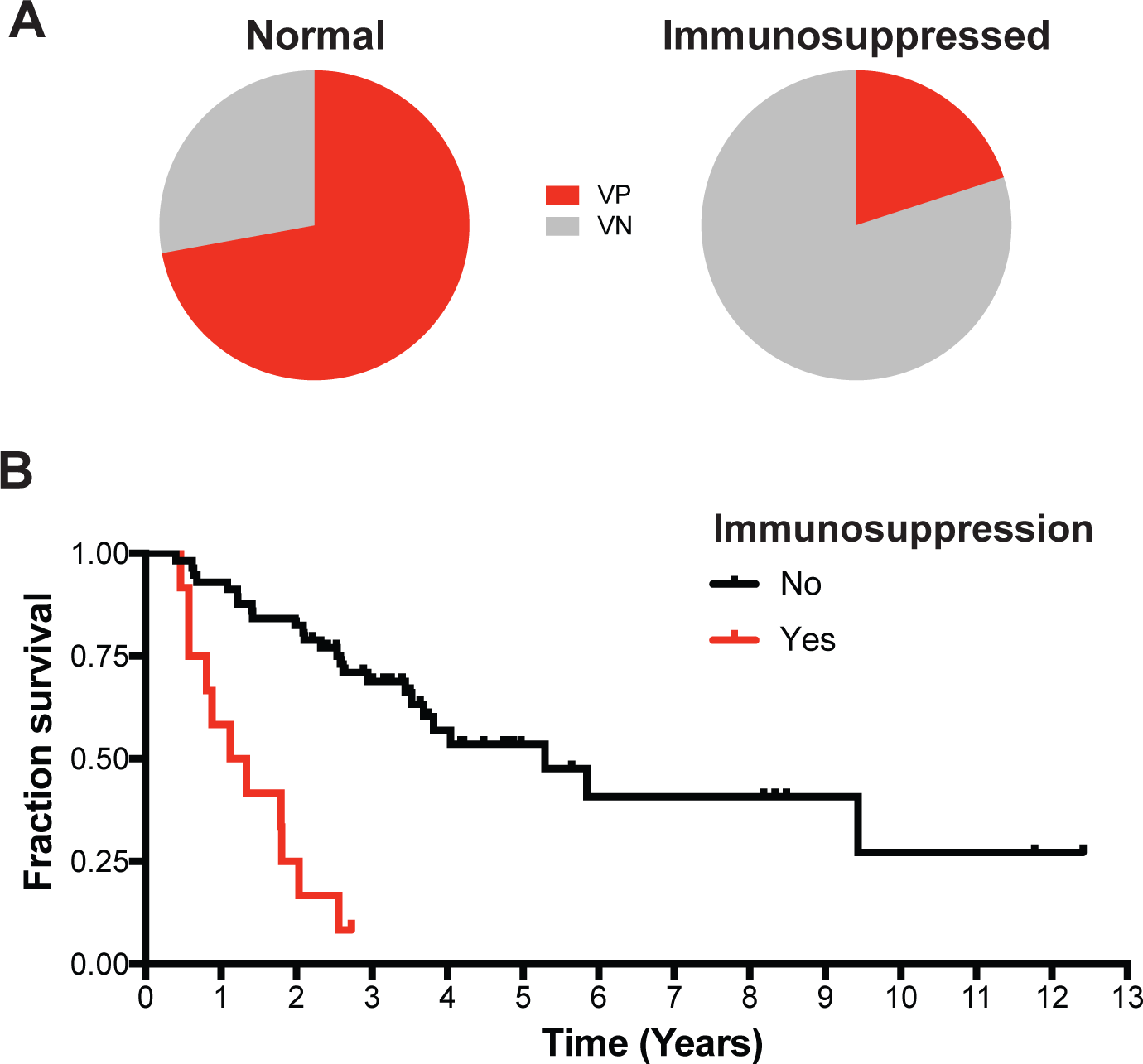
Clinical outcome based on mutation signature, virus status, and immune suppression. A. Pie charts representing the portion of patients that are virus-positive (VP, red) or virus-negative (VN, grey) and immunocompetent or immunosuppressed. B. Kaplan-Meier plot of overall survival of immunocompetent (black) and immunosuppressed (red) MCC patients.

**Table 5.** Association between Patients Characteristics and Immunosuppression using Fisher’s exact test.

| Characteristics | All (N=71) | Immunosuppression |  | Fisher's exact test<br>p-value |
| --- | --- | --- | --- | --- |
|  |  | No (N=61) | Yes (N=10) |  |
| **Gender** |  |  |  |  |
| Female | 31 (44%) | 29 (48%) | 2 (20%) | 0.17 |
| Male | 40 (56%) | 32 (52%) | 8 (80%) |  |
| **Race** |  |  |  |  |
| Black or African American | 2 (3%) | 2 (3%) | 0 (0%) | >0.99 |
| White | 69 (97%) | 59 (97%) | 10 (100%) |  |
| **Age at initial diagnosis, yrs** |  |  |  |  |
| <=70 | 36 (51%) | 32 (52%) | 4 (40%) | 0.51 |
| >70 | 35 (49%) | 29 (48%) | 6 (60%) |  |
| **Initial site** |  |  |  |  |
| Head | 19 (27%) | 14 (23%) | 5 (50%) | 0.39 |
| LE | 15 (21%) | 14 (23%) | 1 (10%) |  |
| Trunk | 23 (32%) | 21 (34%) | 2 (20%) |  |
| UE | 14 (20%) | 12 (20%) | 2 (20%) |  |
| **AJCC stage at initial diagnosis** |  |  |  |  |
| I | 19 (27%) | 16 (26%) | 3 (30%) | 0.96 |
| II | 10 (14%) | 9 (15%) | 1 (10%) |  |
| III | 30 (42%) | 25 (41%) | 5 (50%) |  |
| IV | 12 (17%) | 11 (18%) | 1 (10%) |  |
| **Prior chemotherapy or radiation** |  |  |  |  |
| No | 53 (75%) | 46 (75%) | 7 (70%) | 0.71 |
| Yes | 18 (25%) | 15 (25%) | 3 (30%) |  |
| **UV** |  |  |  |  |
| Negative | 47 (66%) | 45 (74%) | 2 (20%) | <0.01 |
| Positive | 24 (34%) | 16 (26%) | 8 (80%) |  |
| **RB1 status** |  |  |  |  |
| Mutant | 32 (45%) | 24 (39%) | 8 (80%) | 0.04 |
| Wild type | 39 (55%) | 37 (61%) | 2 (20%) |  |
| **TP53 status** |  |  |  |  |
| Mutant | 31 (44%) | 25 (41%) | 6 (60%) | 0.31 |
| Wild type | 40 (56%) | 36 (59%) | 4 (40%) |  |
| **Virus positive or negative** |  |  |  |  |
| VN | 25 (35%) | 17 (28%) | 8 (80%) | <0.01 |
| VP | 46 (65%) | 44 (72%) | 2 (20%) |  |

## Discussion

We undertook this study to develop an assay to more accurately distinguish between virus-positive and virus-negative MCC. We built upon an NGS platform that has been instituted as a routine part of clinical care at the Dana-Farber Cancer Institute, Brigham and Women’s Hospital, and Boston Children’s Hospital. The viral hybrid capture assay, VB2, acquired a high number of MCPyV reads for many samples. Importantly, evidence for specific integration was associated with all cases with a high number of reads (>6,000). Spurious MCPyV reads were also detected in 19 of 20 MCC cases that were deemed to be virus-negative by TMB and UV mutations. There was no evidence for integration in these cases rather these reads could be traced to be extremely low level contamination from MCC037 during library preparation or sequencing. In contrast, true virus-positive MCCs have low TMB with clear assemblies of virus-host junctions with MCC-hallmark deletions in the MCPyV genome.

Integration sites were observed on 12 different chromosomes with the most occurring on chromosome 5. In addition, two fully overlapping integration sites from two different tumors were observed on chromosome 1 separated by only 10-20kb. Based on the clonality of deletions and point mutations in the MCPyV genome, these events most likely occurred before or during integration as was similarly determined from another study on MCC cell lines (34). From previous publications, we can infer that these integration sites likely have amplified adjacent regions of the host genome in a tandem head-to-tail conformation (8). These integration sites frequently are within 50 kb of each other and flank or fully contain enhancer regions of the human genome. These observations are consistent with reports of HPV integration in cervical and head and neck cancers as well as cell lines (35). Particularly, HPV integration events can amplify enhancer regions to high copy number creating super enhancers that can drive expression of adjacent genes and tumor cell survival. This likely represents another important mechanism beyond the expression of the viral tumor antigen in promoting tumorigenesis.

A recurrent amplification of chromosome 6 has previously been observed for MCC; however, this observation predated the discovery of MCPyV and was not associated with morphology or outcome (36). In other cancers, such as basal cell carcinoma and ovarian cancer, this amplification is typically associated with worse outcome (36). Although the chromosome 6 amplification in this study was significantly associated with better overall survival, it was also more frequent in metastasis. This amplification co-occurring with MCPyV may represent a less fatal, but more metastatic subtype of MCC. Additionally, this result could be impacted by the immunotherapy administered to the patients with metastatic MCC.

Unexpectedly, we observed that 8 of 10 cases with immunosuppression were virus-negative MCC. While it was recognized in the early 1990s that individuals with hematologic malignancies that developed MCC had a poor prognosis (37), it was not until 1997 when a direct link between immunosuppression and MCC was postulated (38). At that time, a correlation was noted between medically induced immunosuppression with azathioprine and cyclosporine and the development and rapid spread of MCC. Early reports highlighted a prolonged period of immunosuppression prior to MCC development. The search for a viral pathogen in MCC was initiated because of reports linking MCC with immunosuppression and with HIV-1/AIDS (2).

In the present report, three solid organ transplant recipients, three with chronic auto-immune diseases, and two with hematologic malignancies developed virus-negative MCC. The risk for developing MCC is increased in patients with chronic inflammatory disorders such as rheumatoid arthritis or medically induced immunosuppression for solid organ transplantation (38–41). Skin cancers account for 40-50% of all posttransplant malignancies with squamous cell carcinoma (SCC) and basal cell carcinoma (BCC) comprising 90-95% of these skin cancers (42). Importantly, some therapeutics used in organ transplantation increase risk for developing skin cancers. Azathioprine can sensitize cells to UV-induced damage through the incorporation of a metabolite into DNA that generates reactive oxygen species upon exposure to UV light (43). In patients with rheumatoid arthritis, methotrexate and anti-TNF drugs were associated with an increased risk of non-melanoma skin cancer (44). The increased risk for skin cancers in organ transplant recipients and rheumatoid arthritis is associated with UV-light induced mutagenesis for SCC and BCC. Therefore, the increased risk for UV-induced skin cancers may also extend to virus-negative MCC.

The AIDS-defining malignancies include Kaposi’s sarcoma, driven by human herpes virus 8 (HHV-8) also known as Kaposi’s sarcoma herpes virus (KSHV), non-Hodgkin lymphoma, often triggered by Epstein Barr virus (EBV), and cervical cancer, resulting from human papilloma virus (HPV). Of note, MCC was never categorized as an AIDS-defining malignancy likely due to the rarity of the malignancy even in individuals with profound immunosuppression. Given the association of virus-negative MCC with other forms of immunosuppression observed in this study, it should not be assumed that HIV-1/AIDS associated MCC will be virus-positive MCC.

Despite the significant differences in the TMB between virus-positive and virus-negative MCC, there were few phenotypic differences in the two types of MCC. Based on histopathological features alone, two subtypes of MCC can be recognized: pure neuroendocrine tumors and combined tumors with neuroendocrine and divergent (mainly squamous) differentiation. Most pure tumors are MCPyV-positive and CK20-positive while combined tumors are uniformly MCPyV-negative and occasionally CK-20 negative (9, 45). Virus-negative MCC can also present as pure neuroendocrine type MCC.

While genomic sequencing has revealed that virus-negative MCC has evidence for a high degree of UV damage, this does not exclude a role for UV exposure in the development of virus-positive MCC. The relative lack of UV damaged DNA in virus-positive MCC indicates that the etiologies are clearly different, suggesting that the precursor to virus-negative MCC was a recipient of life-long intense UV exposure while the virus-positive MCC were not exposed to sunlight for the same degree or for as long. It was reported that the early promoter of MCPyV responds to UV exposure and that levels of ST mRNA increased in UV exposed skin from a healthy human volunteer (46). Transient UV exposure could affect the immune response to virus-negative and virus-positive MCC etiology. The effect of UV radiation in the pathogenesis of MCC has been suggested to be more likely a result of immune modulation than direct effects on DNA itself (47).

## Materials and Methods

### Sample collection

DNA was isolated from FFPE sections of MCC tumors corresponding to the 71 individuals summarized in Table 1. In addition, FFPE sections were sectioned for immunohistochemistry with antibodies CM2B4 and Ab3 (17). Sections stained with hematoxylin and eosin were evaluated by synoptic review (32).

### Nucleic acid isolation, library preparation and sequencing

Purified DNA was quantified using a Quant-iT PicoGreen dsDNA assay (ThermoFisher). Library construction was performed using 200 ng of DNA, which was first fragmented to ∼250 bp using a Covaris LE220 Focused ultrasonicator (Covaris, Woburn, MA) followed by size-selected cleanup using Agencourt AMPureXP beads (Beckman Coulter, Inc. Indianapolis, IN) at a 1:1 bead to sample ratio. Fragmented DNA was converted to Illumina libraries using a KAPA HTP library kit using the manufacturer’s recommendations (ThermoFisher). Adapter ligation was done using xGen dual index UMI adapters (IDT, Coralville, IA).

Samples were pooled in equal volume and run on an Illumina MiSeq nano flow cell to quantitate the amount of library based on the number of reads per barcode. All samples yielded sufficient library (> 250 ng) and were taken forward into hybrid capture. Libraries were pooled at equal mass (3 × 17-plex and 1 × 18-plex) to a total of 750 ng. Captures were done using the SureSelect^XT^ Fast target enrichment assay (Agilent, Technologies, Sant Clara, CA) with VB2 with and without supplementation with the OncoPanel (v3) bait set. Captures were sequenced on an Illumina 2500 in rapid run mode (Illumina Inc., San Diego, CA). OncoPanel sequences are available through AACR GENIE (https://www.aacr.org/Research/Research/pages/aacr-project-genie.aspx).

### Sequence alignment and somatic variant calling

Pooled samples were de-multiplexed and sorted using Illumina’s bcl2fastq software (v2.17). Reads were aligned to the reference sequence b37 edition from the Human Genome Reference Consortium as well as viral genomes targeted by the Virus Capture Baitset v2 using bwa mem (http://bio-bwa.sourceforge.net/bwa.shtml) (48). The viral genomes and human genome were combined into one alignment reference so reads could map to the closest matching reference sequence.

Duplicate reads were identified using unique molecular indices (UMIs) and marked using the Picard tools. The alignments were further refined using the Genome Analysis Toolkit (GATK) for localized realignment around indel sites and base quality score recalibration (49, 50).

Mutation analysis for single nucleotide variants (SNV) was performed using MuTect v1.1.4 (CEPH control was used as the “project normal”) and annotated by Variant Effect Predictor v 79 (VEP) (51, 52). We used the SomaticIndelDetector tool that is part of the GATK for indel calling. After initial identification of SNVs and indels by MuTect and GATK respectively, the variants were annotated using OncoAnnotate to determine what genes were impacted and their effect on the amino acid sequence. OncoAnnotate also applied additional filters using the Exome Sequencing Project (ESP) and gnomAD datasets to flag common SNPs.

Variants that affect protein coding regions underwent further filtering/classification based on frequency in the gnomAD, ESP, and COSMIC (version 80) databases. If the frequency of the variant was less than or equal to 1% in all gnomAD and ESP populations, the variant was flagged as “REVIEW_REQUIRED”. If the frequency of the variant was greater than 1% and less than or equal to 10% in all gnomAD and ESP populations and present in “COSMIC” database at least 2 times, the variant was flagged as “REVIEW_REQUIRED”. If the frequency of the variant was between 1% and less than or equal to 10% in all gnomAD and ESP populations and not present in “COSMIC” database at least 2 times, the variant is flagged as “NO_REVIEW_GERMLINE_FILTER”. If the frequency of the variant was greater than 10% in any gnomAD and ESP populations, the variant was flagged as “NO_REVIEW_GERMLINE_FILTER”. Variants with a frequency greater than 10% in any gnomAD or ESP population were considered to be a common SNP irrespective of presence in the COSMIC database.

Variants in the viral genomes were called using samtools mpileup and bcftools from the aligned bam files. Called variants were filtered to have a minimum coverage of 5 reads and minimum allele frequency of 1% of total reads covering that base in a single sample. Variants were annotated based on the NC_010277.2 reference sequence in GenBank using SnpEff (PMID: 22728672).

### Recurrent copy number analysis

Copy number variant calling was performed using a combination of VisCap Cancer and CNVkit as previously described (23)(53). All resulting gene copy number variants from all patients were compared against each other with UV status and significant mutual exclusivity/co-occurrence was calculated using Fisher’s exact test corrected by FDR for multiple comparisons in the R statistical environment. Using the network and iGraphs packages the significantly co-occurrent variants were clustered into networks. The genes belonging to each distinct network cluster with more than 5 member genes were then labeled and extracted. Using these gene lists as cluster definitions each patient was evaluated for presence or absence of each CNV cluster. Presence of a CNV cluster was determined if more than 50% of the member genes of that cluster were modified in the same patient. Code is available from https://github.com/gstarrett/oncovirus_tools.

### Viral integration analysis

A custom perl script was written to extract, assemble, annotate, and visualize viral reads and determine viral integration sites. Viral reads and their mates were first identified and extracted by those that have at least one mate map to the viral genome. Additional reads containing viral sequence were identified by a bloom filter constructed of unique, overlapping 31bp k-mers of the MCPyV genome (54). The human genome positions for any read with a mate mapping to the viral genome were output into a bed file and the orientation of viral and human pairs was stored to accurately deconvolute overlapping integration sites. This bed file was then merged down into overlapping ranges based on orientation counting the number of reads overlapping that range. Skewdness in coverage of integration junctions was calculated by the difference in the fraction of virus-host read pairs overlapping the first and second halves of the aforementioned ranges. This skewdness value was used to determine the orientation of the viral-host junction (i.e. positive values, junction is on the 3’ end of the range; negative values, junction is on the 5’ end of the range), which was validated from the results of de novo assembly. Integrated viral genomes were assembled from extracted reads using SPAdes with default parameters (55). The assembly graphs from SPAdes were annotated using blastn against hg19 and the MCPyV reference genome with an e-value cutoff of 1×10^−10^. Annotated assembly graphs were visualized using the ggraph R package. Code is available from https://github.com/gstarrett/oncovirus_tools.

Integrations sites confirmed by reference guided alignment and assembly data were analyzed for stretches of microhomology between the human and viral genomes by selecting 10bp upstream and downstream of the integration junction on the viral and human genomes. Within these sequences stretches of identical sequence at the same position longer than two base pairs were counted. Overall homology between the sequences was calculated by Levenshtein distance (). Three integration junctions with indeterminate DNA sequence ranging from 1-25bp inserted between viral and human DNA were excluded from analysis. Expected microhomology was calculated by randomly selecting 1000 20bp pairs of non-N containing sequence from the human and MCPyV genomes.

Integration site proximity to repeat elements were determined using bedtools closest and repeatmasker annotations acquired from the UCSC genome browser (56). Expected frequency of integration near repeat elements were determined by randomly selecting 1000 sites in the human genome. Sites within 2kb of a repeat element were counted as close proximity.

### Statistics

The association between relapse and genomic characteristics are tested with Fisher’s exact test. Overall survival (OS) is defined as the time from initial diagnosis to death, and patients who did not die are censored at the date last follow-up date. The 95% confidence intervals of the median OS times are estimated using log(-log(OS)) methodology. Multivariable Cox regression model is built to investigate the association between virus status and overall survival adjusted by other covariates. And model is fit using backward, forward, and stepwise selection to assess model consistency. Hazard ratios are presented with 95% Wald confidence intervals. Statistical significance is defined as p ≤ 0.05.

Associations between recurrent CNV, TMB, or viral copies and overall survival were calculated and graphed using Graphpad Prism 7. Fisher’s exact test and Kaplan-Meier curves were computed with the R statistical environment. Significant enrichment of microhomology and repeat elements at integrations sites was determined using Fisher’s exact test between observed and expected events.

### Human subjects

This study was conducted according to Declaration of Helsinki principles and approved by the Dana-Farber Cancer Institute institutional review board. Written informed consent was received from participants prior to inclusion in the study.

## Supporting information

Supplement table of contents

## Author Contributions

Conceptualization: G.J.S., L.E.M., J.A.D.

Methodology: G.J.S., T.C., L.E.M., J.A.D.

Software: G.J.S., A.R.T.

Validation: G.J.S., J.C.

Formal Analysis: G.J.S., T.C., J.N., R.T.B., J.A.D.

Investigation: M.T., C.M., J.C., A.R.T, W.P., M.S., A.P., F.K., J.A.D.

Resources: M.T., C.P., G.R., A.G.-H., L.E.M., J.A.D.

Data Curation: G.J.S., T.C., C.P. A.R.T., L.E.M., J.A.D.

Writing-Original Draft: G.J.S., T.C., A.R.T., J.A.D.

Writing-Reviewing & Editing: G.J.S., L.E.M., J.A.D.

Visualization: G.J.S.

Supervision: A.R.T., A.G.-H., L.E.M., J.A.D.

Project Administration: A.G.-H., L.E.M., J.A.D.

Funding Acquisition: J.A.D.

## Acknowledgements

This work was supported in part by US Public Health Service grants (report.nih.gov) R01CA63113, R01CA173023 and P01CA203655 to JAD. JAD has received honoraria for participation in advisory board from Merck & Co., Inc. JAD has received research funding from Constellation Pharmaceuticals, Inc. Salary support for GJS comes from the NCI’s cancer research training award. We thank Christopher B. Buck (NCI) for helpful comments and discussion.

The authors acknowledge the DFCI Oncology Data Retrieval System (OncDRS) for the aggregation, management, and delivery of the clinical and operational research data used in this project. The content is solely the responsibility of the authors.

