## Supplement table of contents for "Clinical and molecular characterization of virus-positive and virus-negative Merkel cell carcinoma"

**Supplemental Information**

Clinical and molecular characterization of virus-positive and virus-negative Merkel cell carcinoma with implications for overall survival

**Authors**:

Gabriel J. Starrett[1], Manisha Thakuria[2,3], Tianqi Chen[4], Christina Marcelus[5], Jingwei Cheng[5,6], Jason Nomburg[5], Aaron R. Thorner[7], Michael K. Slevin[7], Winslow Powers[7], Robert T. Burns[7], Caitlin Perry[4], Adriano Piris[2], Frank C. Kuo[8], Guilherme Rabinowits[3,5,9], Anita Giobbie-Hurder[4], Laura E. MacConaill[7,8], James A. DeCaprio[3,5,6,10]

1. Laboratory of Cellular Oncology, CCR, NCI, NIH, Bethesda, MD

2. Department of Dermatology, Brigham and Women's Hospital, Harvard Medical School, Boston, MA

3. Merkel Cell Carcinoma Center of Excellence, Dana-Farber/Brigham Cancer Center, Boston, MA

4. Department of Data Sciences, Dana-Farber Cancer Institute, Boston, MA

5. Department of Medical Oncology, Dana-Farber Cancer Institute, Boston, MA

6. Department of Medicine, Brigham and Women's Hospital, Harvard Medical School, Boston, MA

7. Center for Cancer Genome Discovery, Dana-Farber Cancer Institute, Boston, MA

8. Department of Pathology, Brigham and Women's Hospital, Harvard Medical School, Boston, MA

9. Current address: Miami Cancer Institute, Miami, FL

10. Corresponding author

**Table of Contents:**

Figure S1. Oncoprint for all genes in this study
Figure S2. Network graph for recurrent CNVs
Figure S3. CNV frequency by cluster for all patients
Figure S4. Assembly graphs for all integration events

Table S1. SNV data for all patients
Table S2. CNV data for all patients
Table S3. CNV cluster definitions
Table S4. Synoptic review of dermatopathology
Table S5. Association between Relapse and Genomic Sequencing (N=52)

### Figure Legends

**Figure S1. Oncoprint for all genes in this study**. Sample are in order of descending TMB and genes are in order of highest point mutations to least.

**Figure S2. Network graph for recurrent CNVs.** Network visualization produced by igraph for genes significantly co-mutated in MCC.

**Figure S3. CNV frequency by cluster for all patients**. Cluster number is shown in grey bars above the bar plots representing amplification/sgains (red) and deletions/losses (blue). Below the bar plots is a heat map of all CNVs (genes, x-axis) (amplifications/gains, red; deletions/losses, blue; no change, grey) across all samples (y-axis) annotated by cluster and chromosome. On the left side pRB, p53 shown in grey and black for 1 or 2 copy loss/mutant, respectively. Presence of UV mutations are shown in black.

**Figure S4. Assembly graphs for all integration events**. 28 assembly graphs annotated by MCPyV genome position (colors labeling each segment are under the header “as.factor(V9)”). The “V3” variable represents the coverage of each contig as determined by SPAdes and contigs are scaled to reflect this value.
