## Supplementary figures and images for "Clinical and molecular characterization of virus-positive and virus-negative Merkel cell carcinoma"

### Figure S1

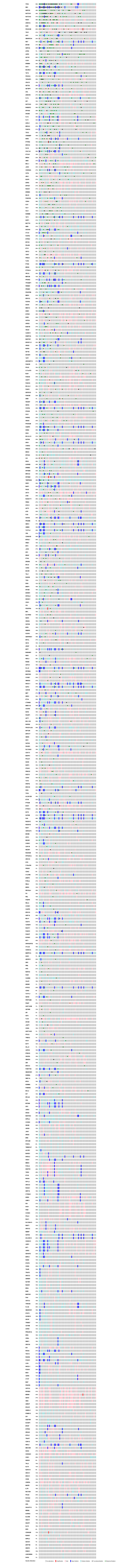

### Figure S2

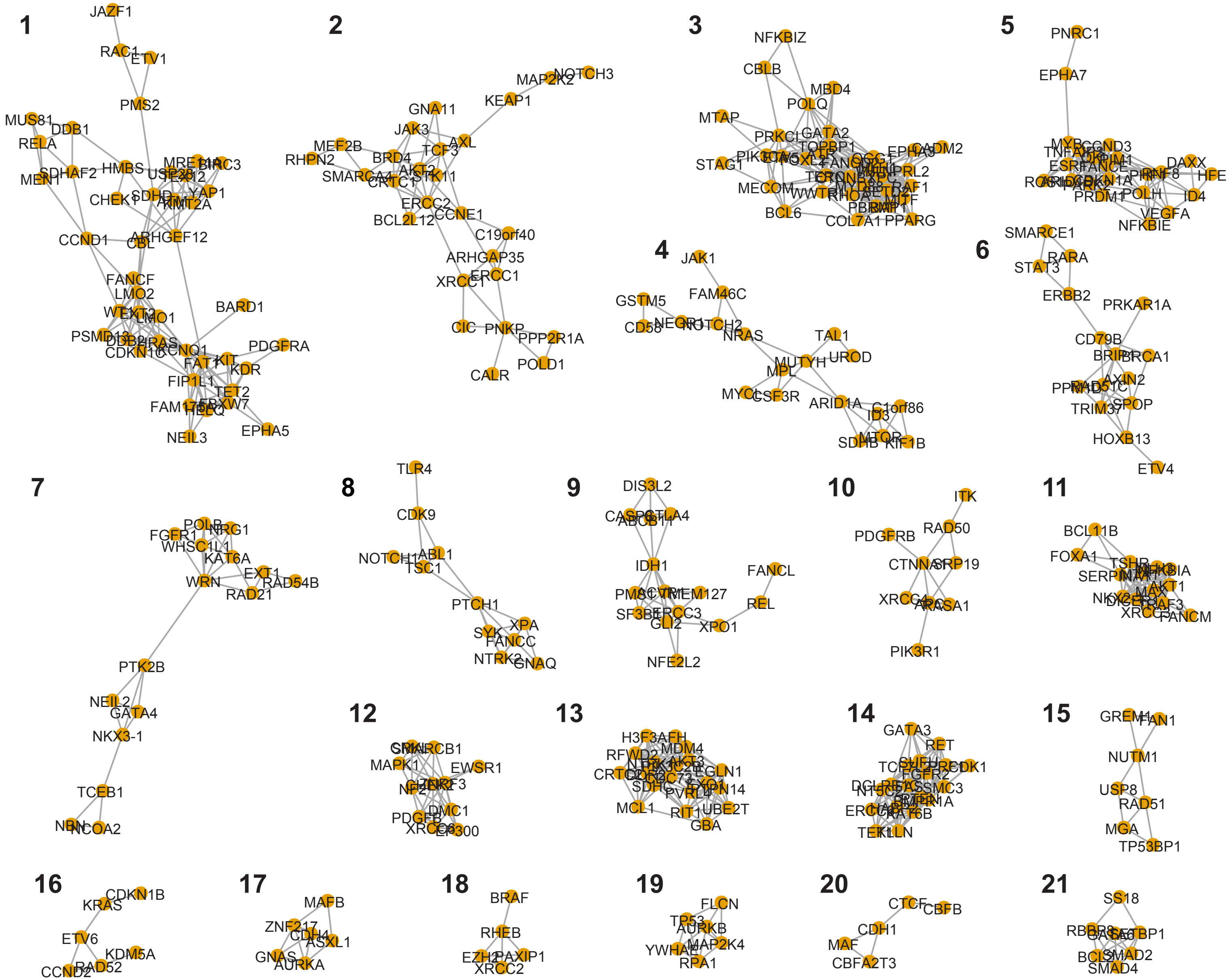

### Figure S3

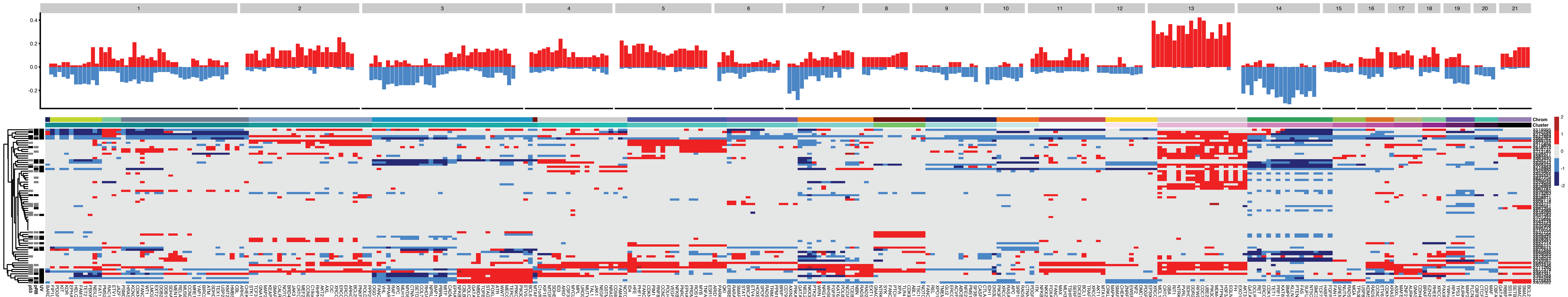
