## Supplementary material for "Clinical and molecular characterization of virus-positive and virus-negative Merkel cell carcinoma": Figure S4

MCC001.pdf

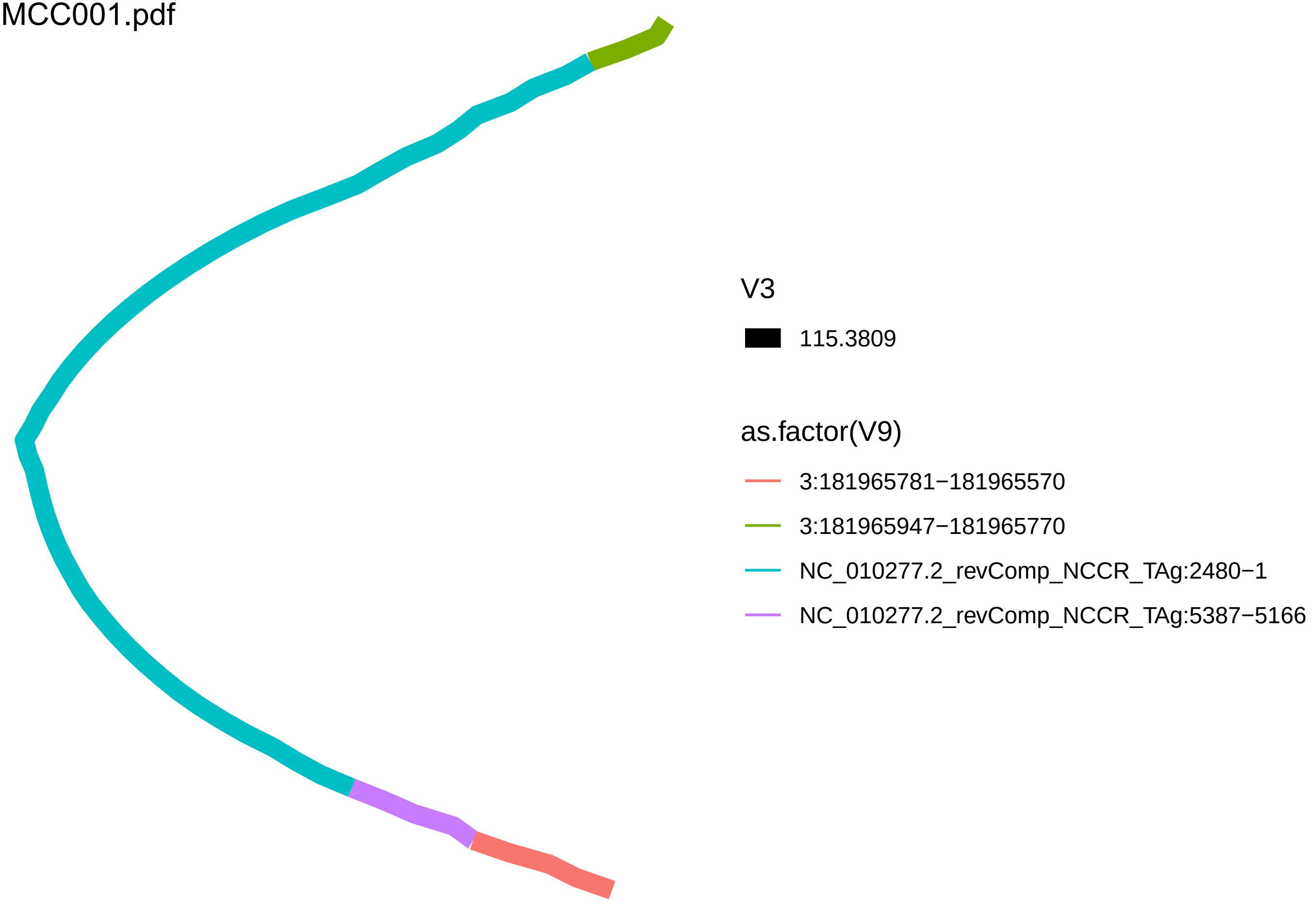

V3
115.3809
as.factor(V9)
3:181965781−181965570
3:181965947−181965770
NC_010277.2_revComp_NCCR_TAg:2480−1
NC_010277.2_revComp_NCCR_TAg:5387−5166

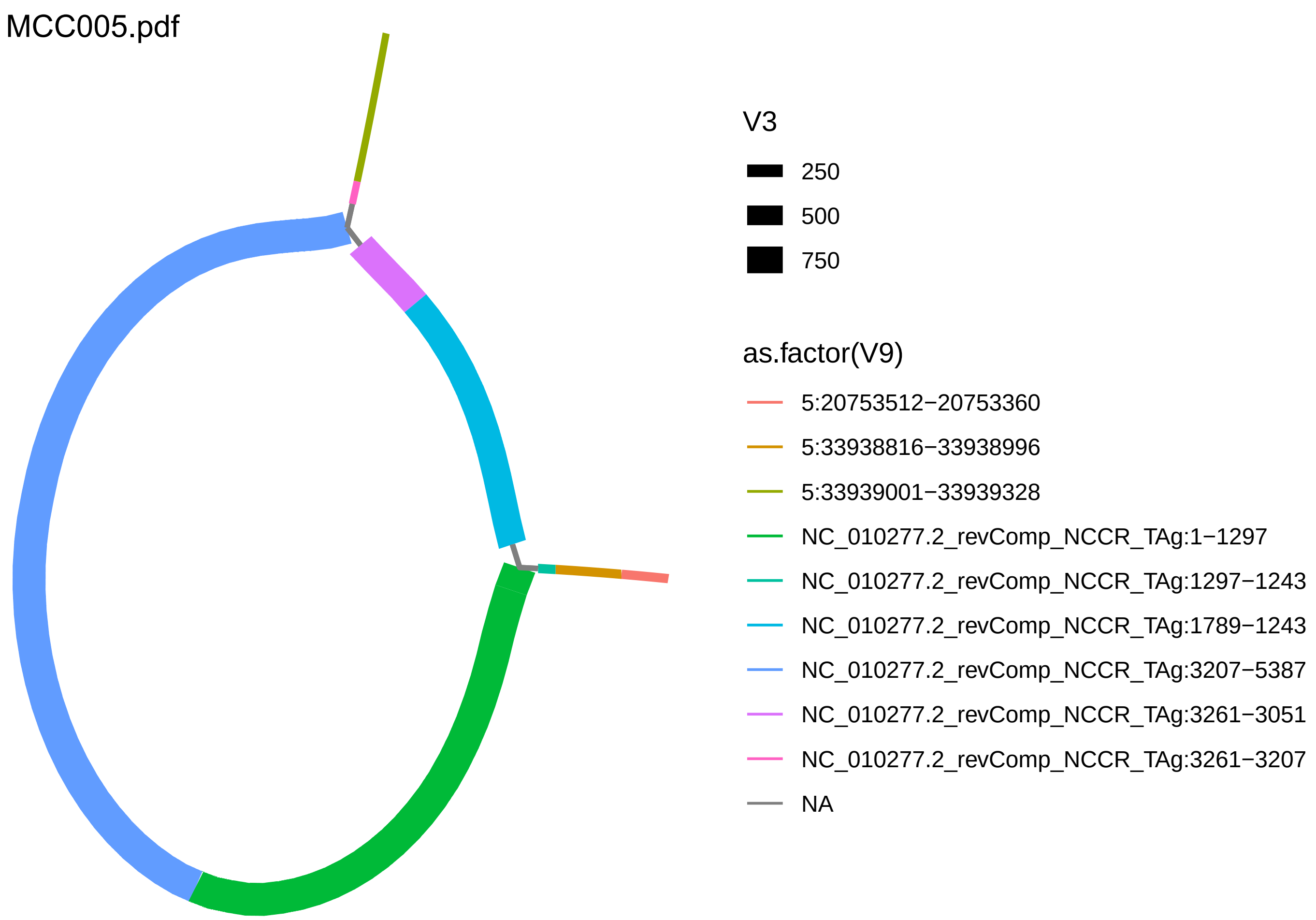
MCC005.pdf
V3
250
500
750
as.factor(V9)
5:20753512−20753360
5:33938816−33938996
5:33939001−33939328
NC_010277.2_revComp_NCCR_TAg:1−1297
NC_010277.2_revComp_NCCR_TAg:1297−1243
NC_010277.2_revComp_NCCR_TAg:1789−1243
NC_010277.2_revComp_NCCR_TAg:3207−5387
NC_010277.2_revComp_NCCR_TAg:3261−3051
NC_010277.2_revComp_NCCR_TAg:3261−3207
NA

MCC006.pdf

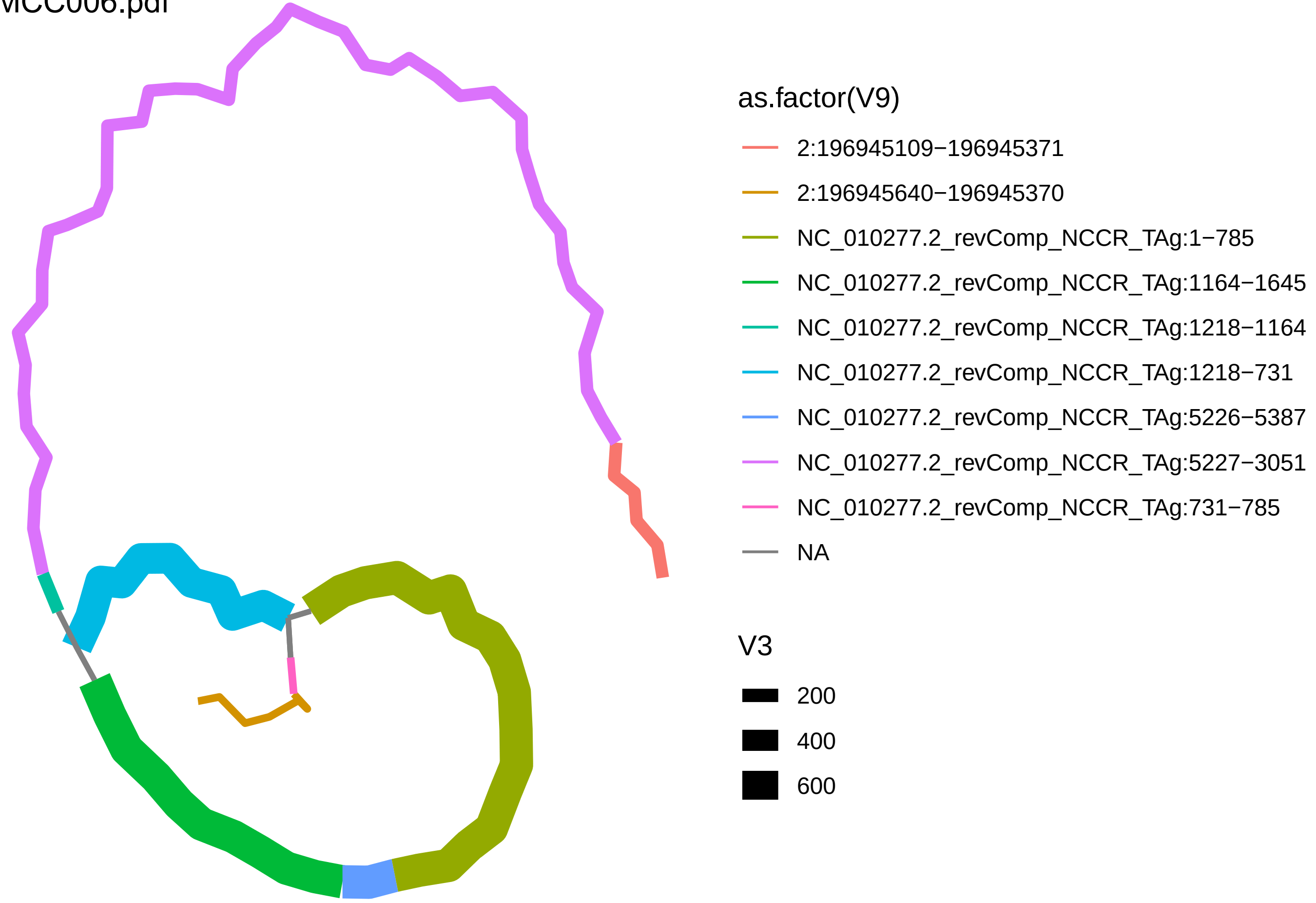

as.factor(V9)
2:196945109−196945371
2:196945640−196945370
NC_010277.2_revComp_NCCR_TAg:1−785
NC_010277.2_revComp_NCCR_TAg:1164−1645
NC_010277.2_revComp_NCCR_TAg:1218−1164
NC_010277.2_revComp_NCCR_TAg:1218−731
NC_010277.2_revComp_NCCR_TAg:5226−5387
NC_010277.2_revComp_NCCR_TAg:5227−3051
NC_010277.2_revComp_NCCR_TAg:731−785
NA
V3
200
400
600

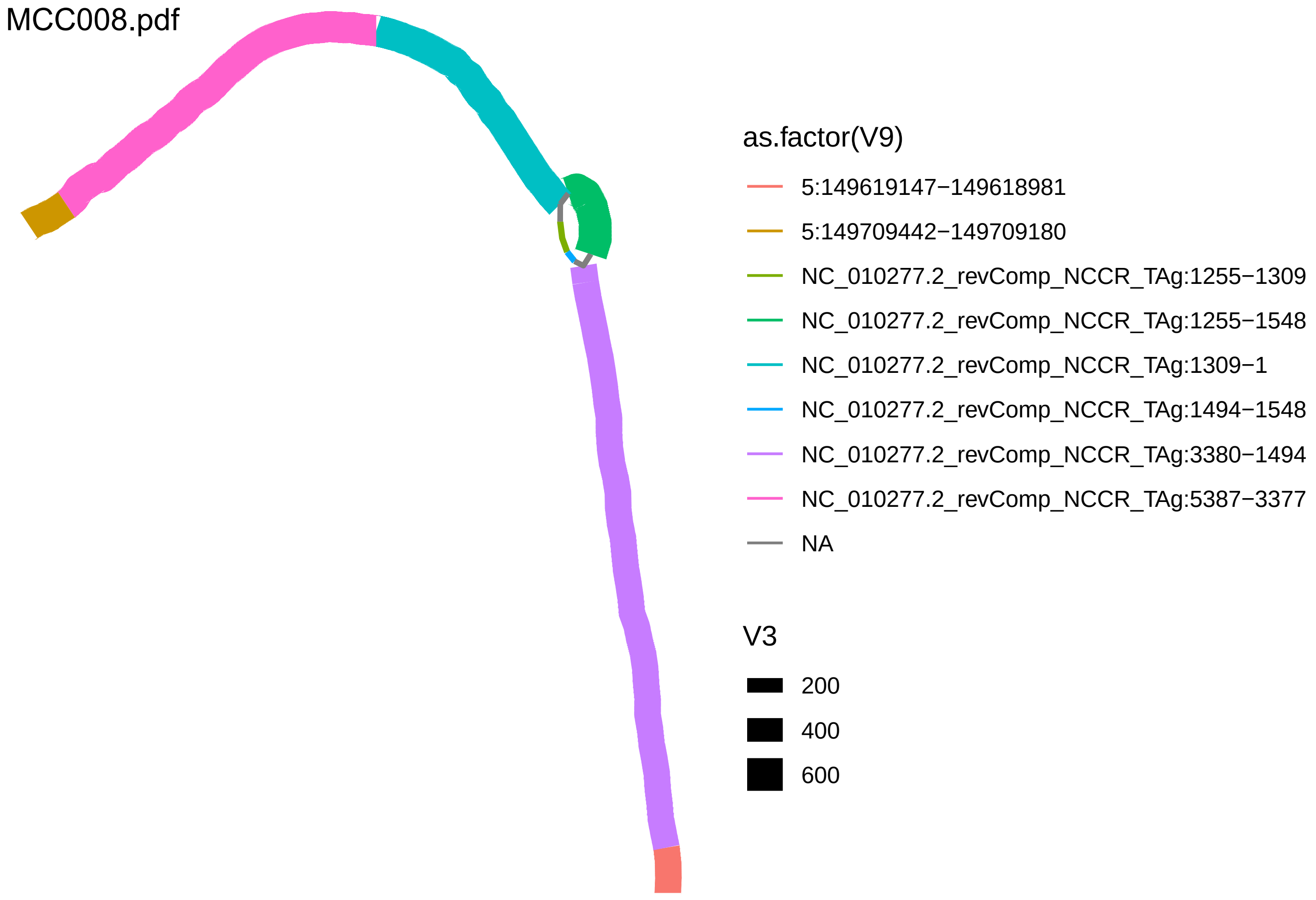
MCC008.pdf
as.factor(V9)
5:149619147−149618981
5:149709442−149709180
NC_010277.2_revComp_NCCR_TAg:1255−1309
NC_010277.2_revComp_NCCR_TAg:1255−1548
NC_010277.2_revComp_NCCR_TAg:1309−1
NC_010277.2_revComp_NCCR_TAg:1494−1548
NC_010277.2_revComp_NCCR_TAg:3380−1494
NC_010277.2_revComp_NCCR_TAg:5387−3377
NA
V3
200
400
600

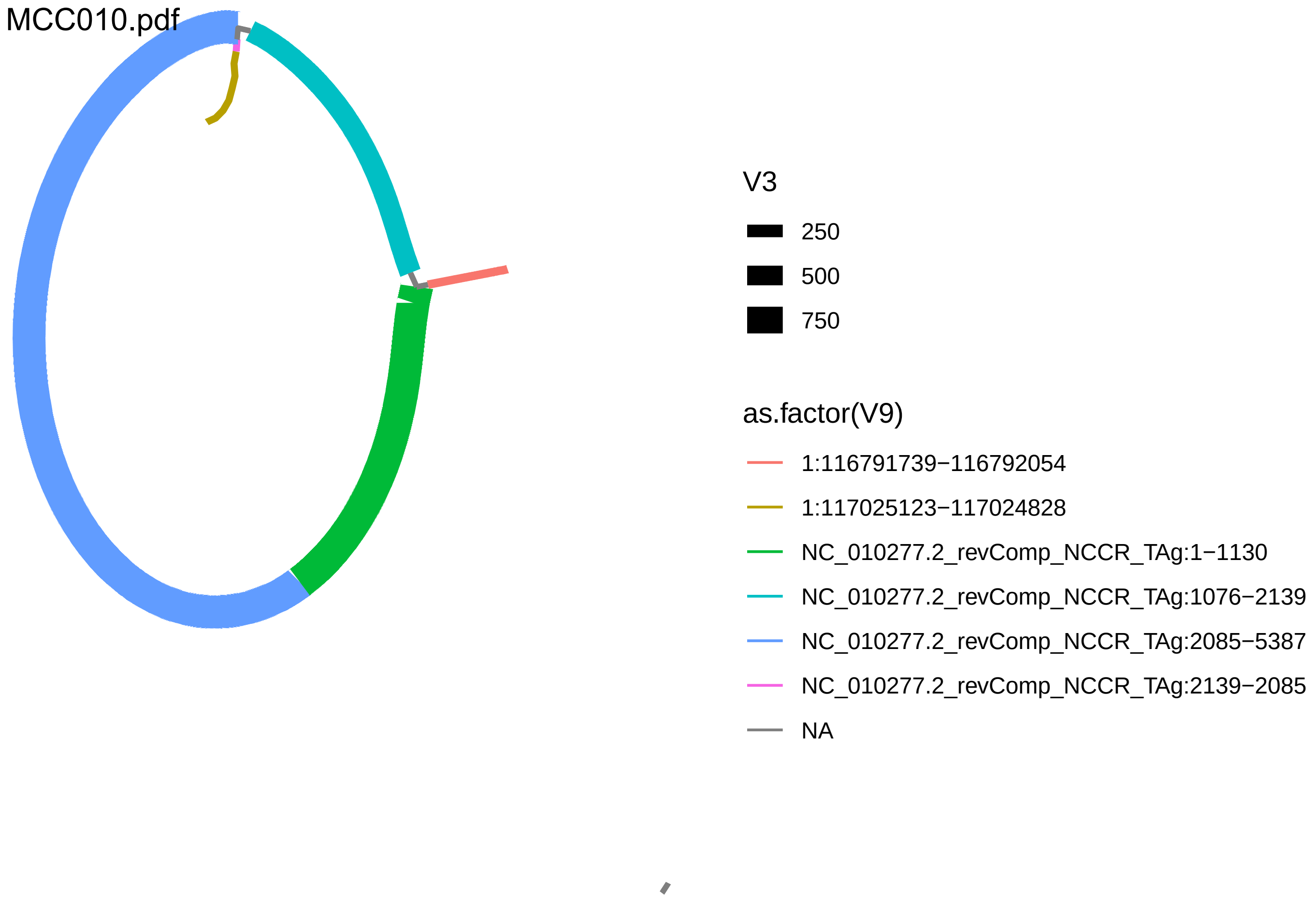

MCC010.pdf
V3
250
500
750
as.factor(V9)
1:116791739−116792054
1:117025123−117024828
NC_010277.2_revComp_NCCR_TAg:1−1130
NC_010277.2_revComp_NCCR_TAg:1076−2139
NC_010277.2_revComp_NCCR_TAg:2085−5387
NC_010277.2_revComp_NCCR_TAg:2139−2085
NA

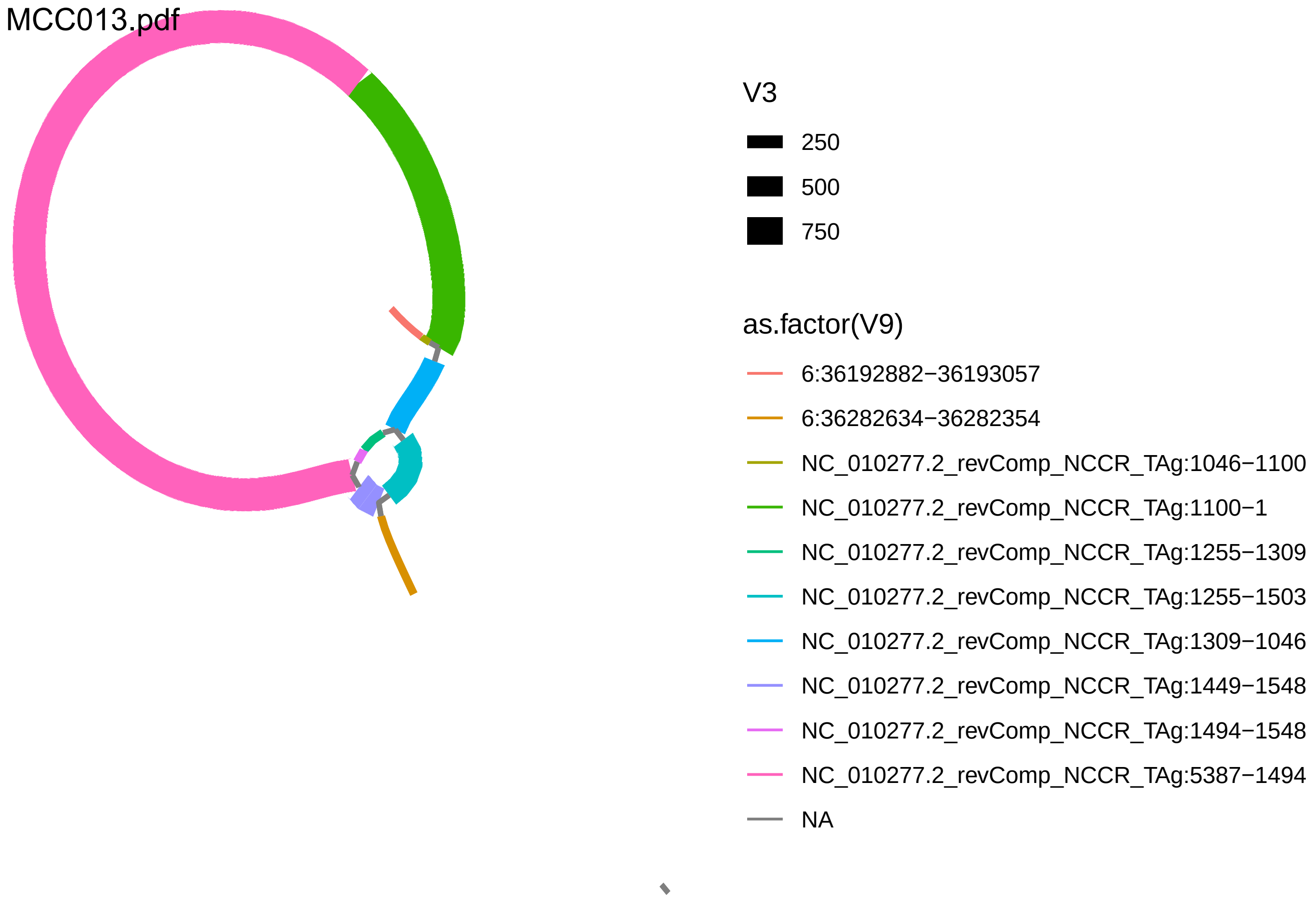
MCC013.pdf
V3
250
500
750
as.factor(V9)
6:36192882−36193057
6:36282634−36282354
NC_010277.2_revComp_NCCR_TAg:1046−1100
NC_010277.2_revComp_NCCR_TAg:1100−1
NC_010277.2_revComp_NCCR_TAg:1255−1309
NC_010277.2_revComp_NCCR_TAg:1255−1503
NC_010277.2_revComp_NCCR_TAg:1309−1046
NC_010277.2_revComp_NCCR_TAg:1449−1548
NC_010277.2_revComp_NCCR_TAg:1494−1548
NC_010277.2_revComp_NCCR_TAg:5387−1494
NA

MCC014.pdf

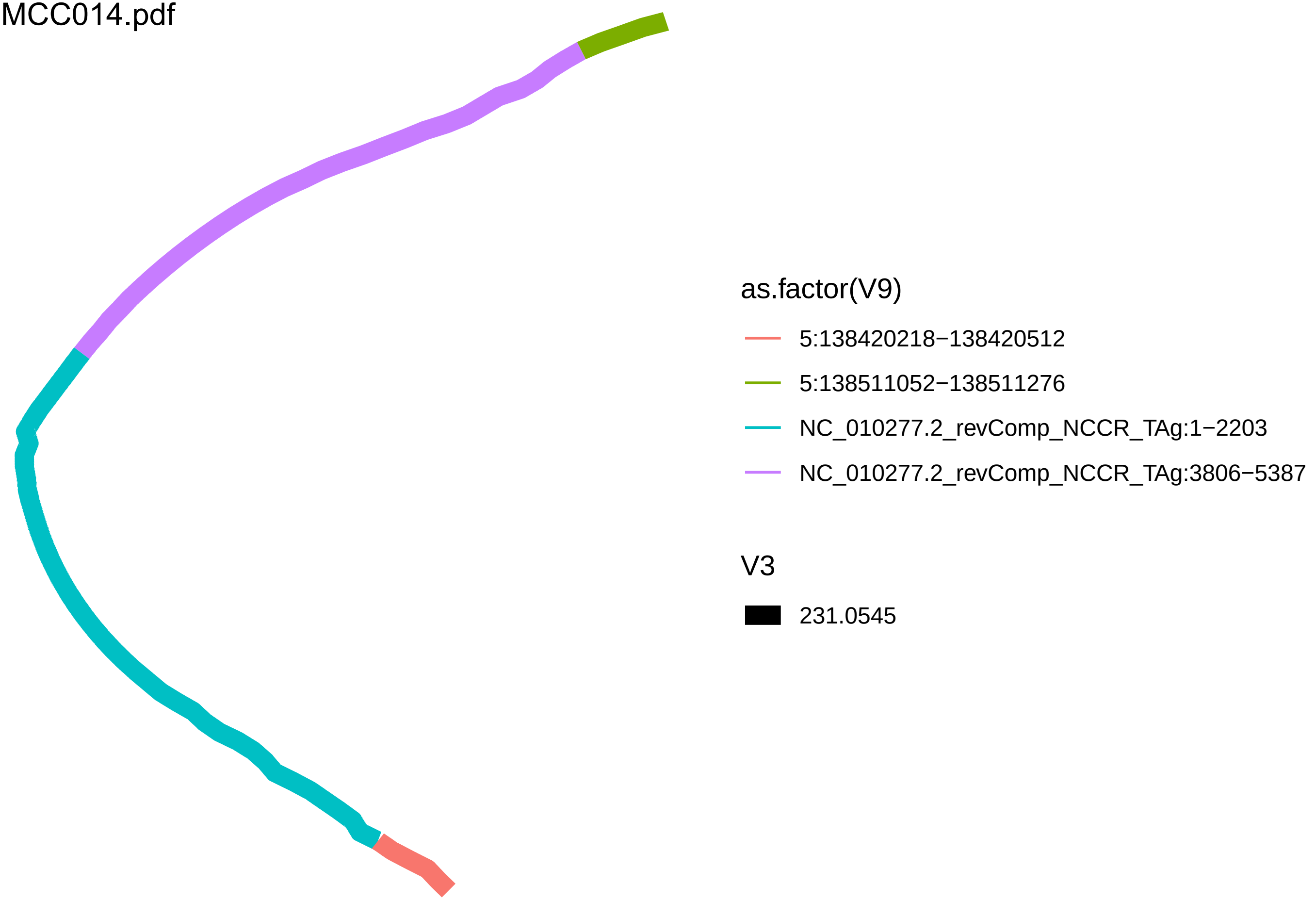

as.factor(V9)
5:138420218−138420512
5:138511052−138511276
NC_010277.2_revComp_NCCR_TAg:1−2203
NC_010277.2_revComp_NCCR_TAg:3806−5387
V3
231.0545

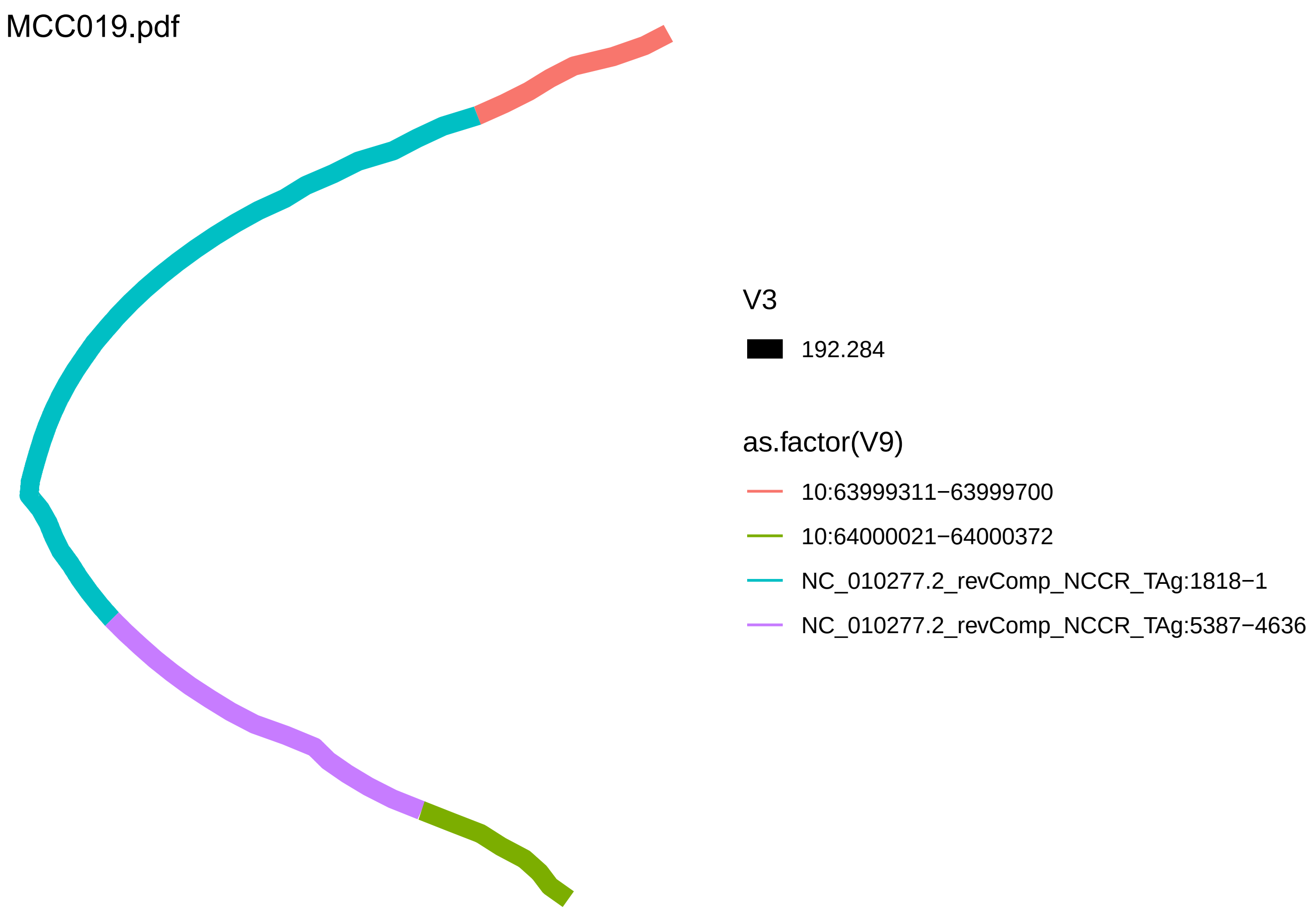
MCC019.pdf
V3
192.284
as.factor(V9)
10:63999311−63999700
10:64000021−64000372
NC_010277.2_revComp_NCCR_TAg:1818−1
NC_010277.2_revComp_NCCR_TAg:5387−4636

MCC022.pdf

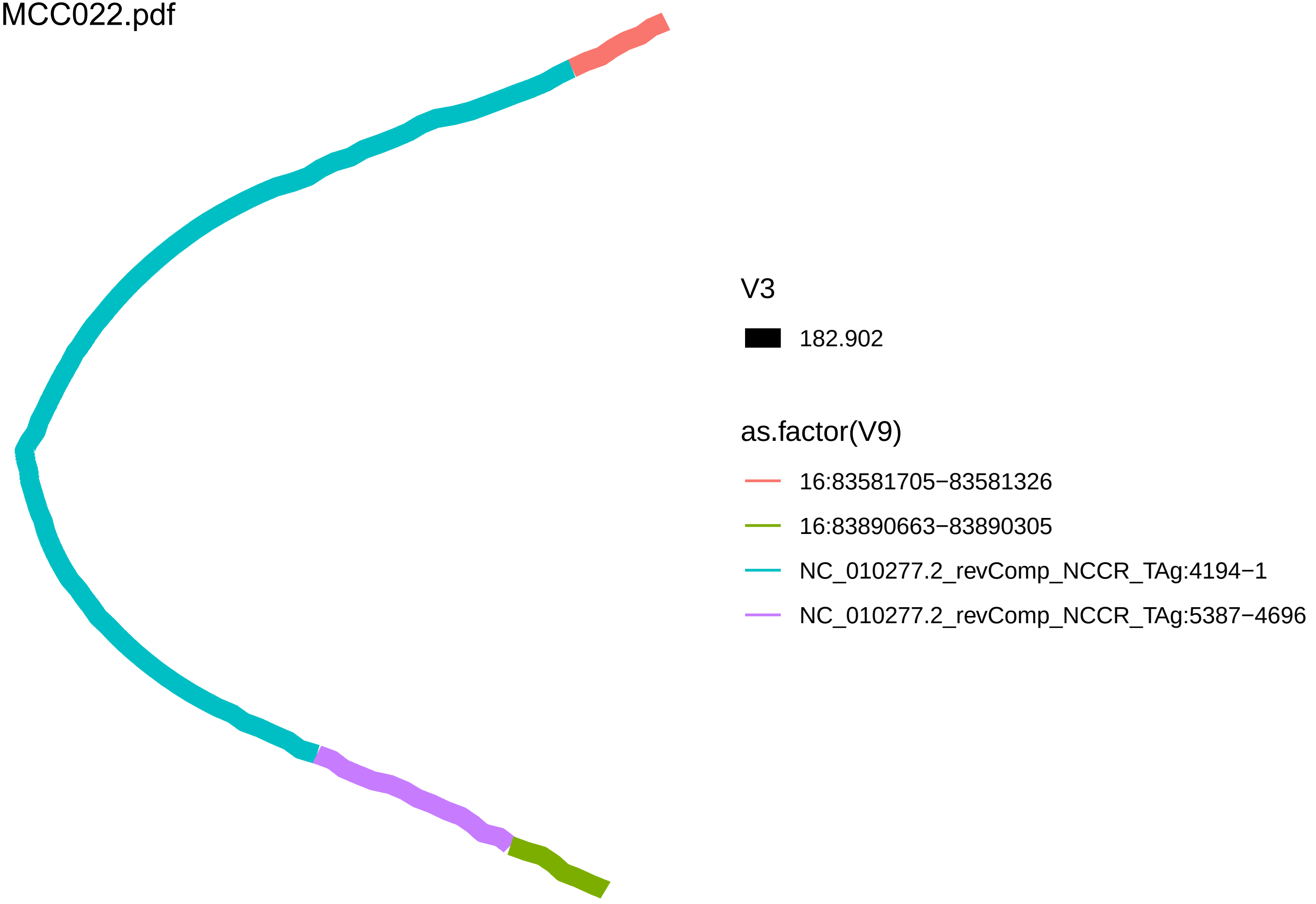

V3
182.902
as.factor(V9)
16:83581705−83581326
16:83890663−83890305
NC_010277.2_revComp_NCCR_TAg:4194−1
NC_010277.2_revComp_NCCR_TAg:5387−4696

MCC024.pdf

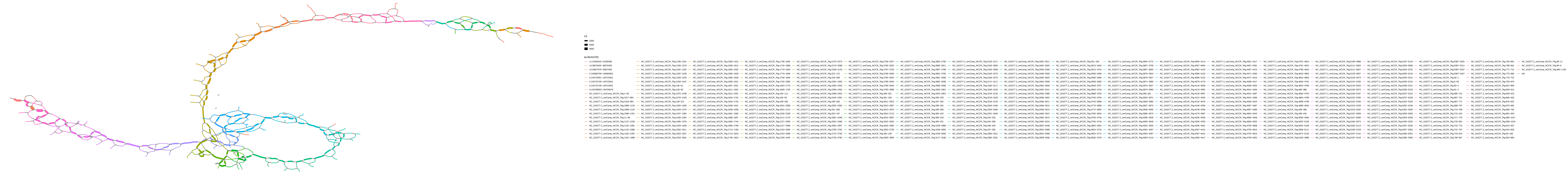

V3
3000
6000
9000
as.factor(V9)
11:31565332−31565386
14:38576492−38576400
14:45827976−45827930
5:145660706−145660952
5:145722651−145722831
5:145722705−145722641
5:145722705−145722651
5:145766003−145765679
NC_010277.2_revComp_NCCR_TAg:1−56
NC_010277.2_revComp_NCCR_TAg:1238−1291
NC_010277.2_revComp_NCCR_TAg:1238−1379
NC_010277.2_revComp_NCCR_TAg:1291−1182
NC_010277.2_revComp_NCCR_TAg:1292−1238
NC_010277.2_revComp_NCCR_TAg:1328−1429
NC_010277.2_revComp_NCCR_TAg:1328−1437
NC_010277.2_revComp_NCCR_TAg:136−61
NC_010277.2_revComp_NCCR_TAg:136−93
NC_010277.2_revComp_NCCR_TAg:1375−1438
NC_010277.2_revComp_NCCR_TAg:1556−1431
NC_010277.2_revComp_NCCR_TAg:1558−1612
NC_010277.2_revComp_NCCR_TAg:1583−1502
NC_010277.2_revComp_NCCR_TAg:1585−1529
NC_010277.2_revComp_NCCR_TAg:1585−1639
NC_010277.2_revComp_NCCR_TAg:1605−1531
NC_010277.2_revComp_NCCR_TAg:1607−1551
NC_010277.2_revComp_NCCR_TAg:1611−1502
NC_010277.2_revComp_NCCR_TAg:1612−1551

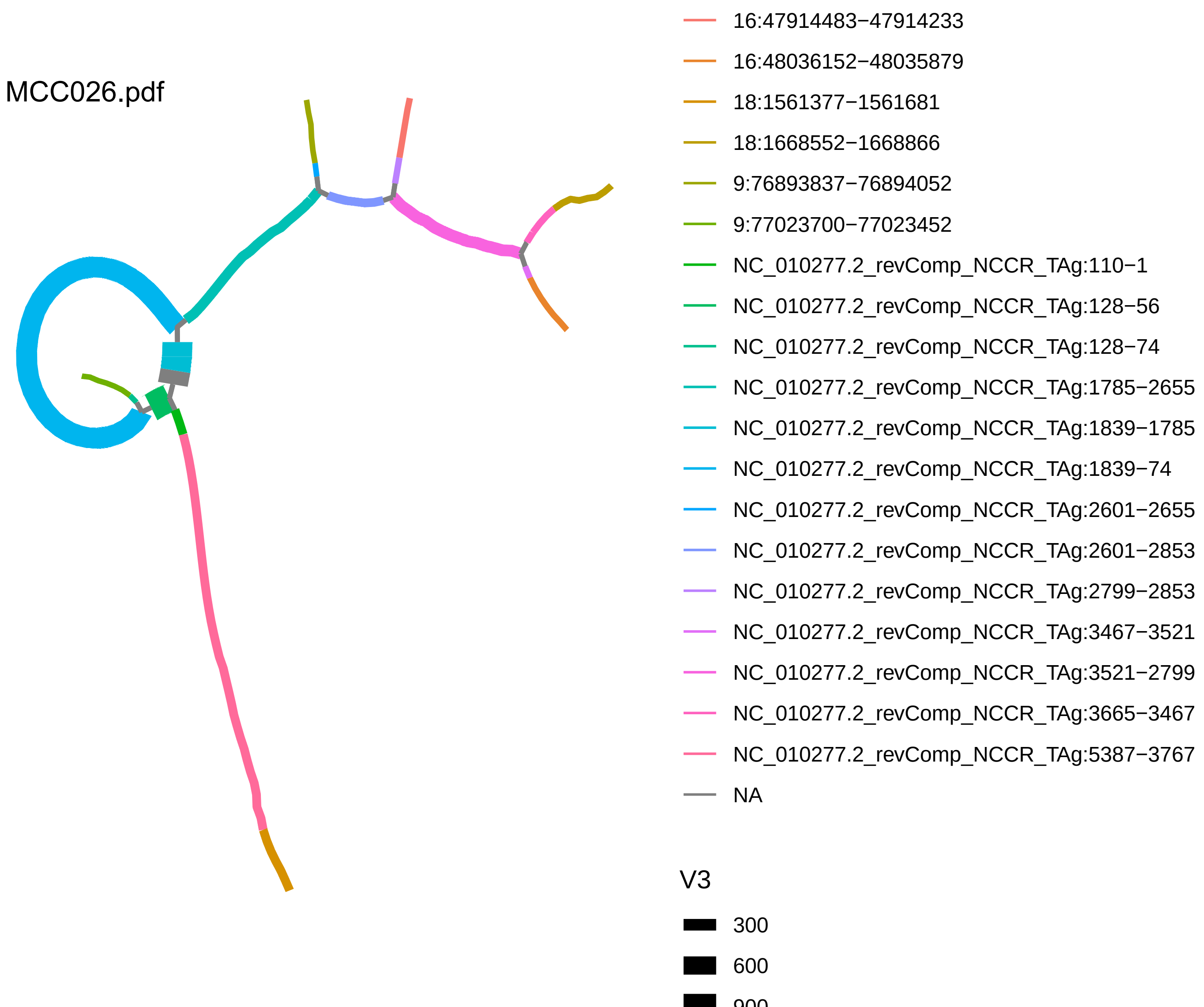

MCC026.pdf
16:47914483−47914233
16:48036152−48035879
18:1561377−1561681
18:1668552−1668866
9:76893837−76894052
9:77023700−77023452
NC_010277.2_revComp_NCCR_TAg:110−1
NC_010277.2_revComp_NCCR_TAg:128−56
NC_010277.2_revComp_NCCR_TAg:128−74
NC_010277.2_revComp_NCCR_TAg:1785−2655
NC_010277.2_revComp_NCCR_TAg:1839−1785
NC_010277.2_revComp_NCCR_TAg:1839−74
NC_010277.2_revComp_NCCR_TAg:2601−2655
NC_010277.2_revComp_NCCR_TAg:2601−2853
NC_010277.2_revComp_NCCR_TAg:2799−2853
NC_010277.2_revComp_NCCR_TAg:3467−3521
NC_010277.2_revComp_NCCR_TAg:3521−2799
NC_010277.2_revComp_NCCR_TAg:3665−3467
NC_010277.2_revComp_NCCR_TAg:5387−3767
NA
V3
300
600
900

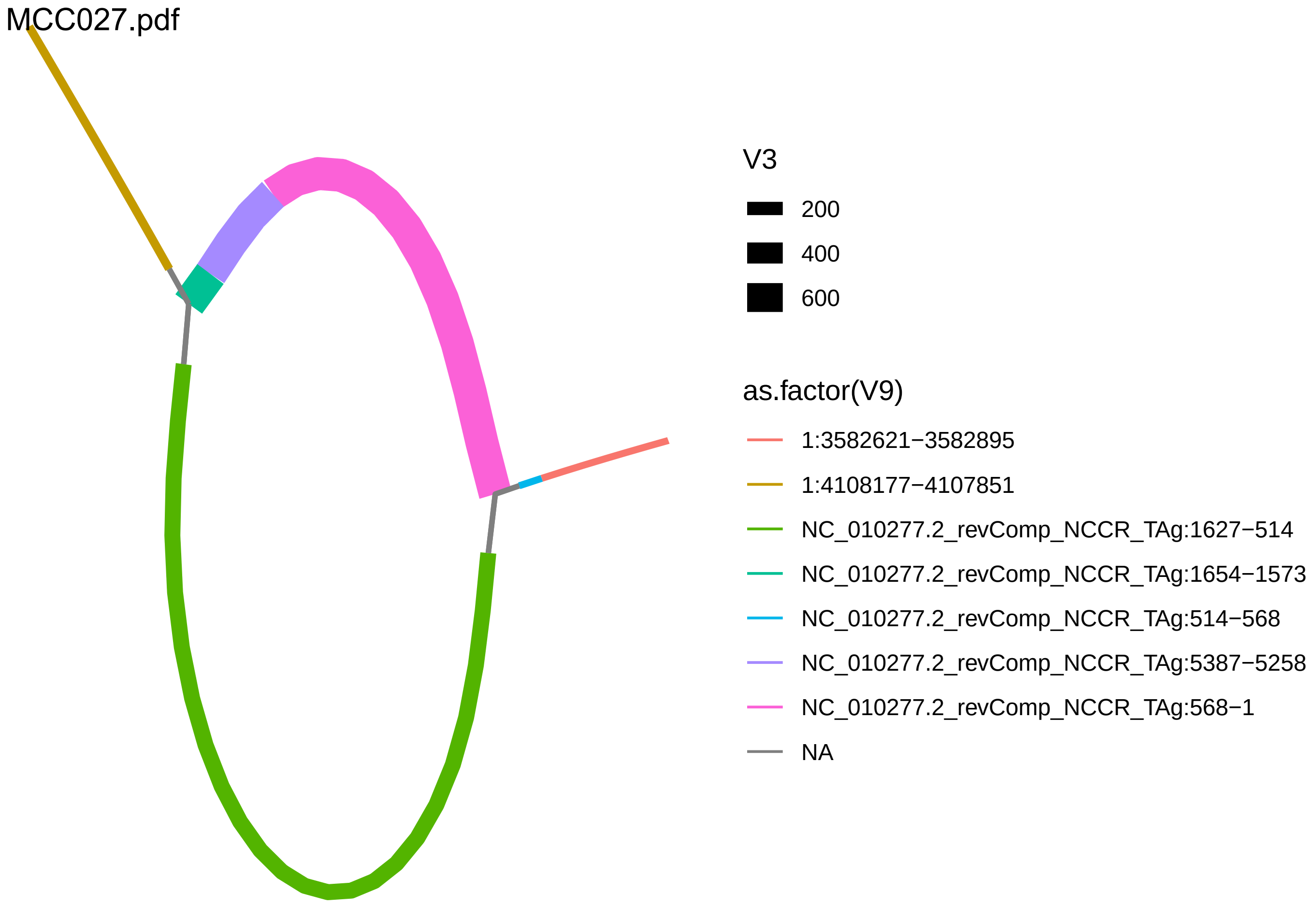
MCC027.pdf
V3
200
400
600
as.factor(V9)
1:3582621−3582895
1:4108177−4107851
NC_010277.2_revComp_NCCR_TAg:1627−514
NC_010277.2_revComp_NCCR_TAg:1654−1573
NC_010277.2_revComp_NCCR_TAg:514−568
NC_010277.2_revComp_NCCR_TAg:5387−5258
NC_010277.2_revComp_NCCR_TAg:568−1
NA

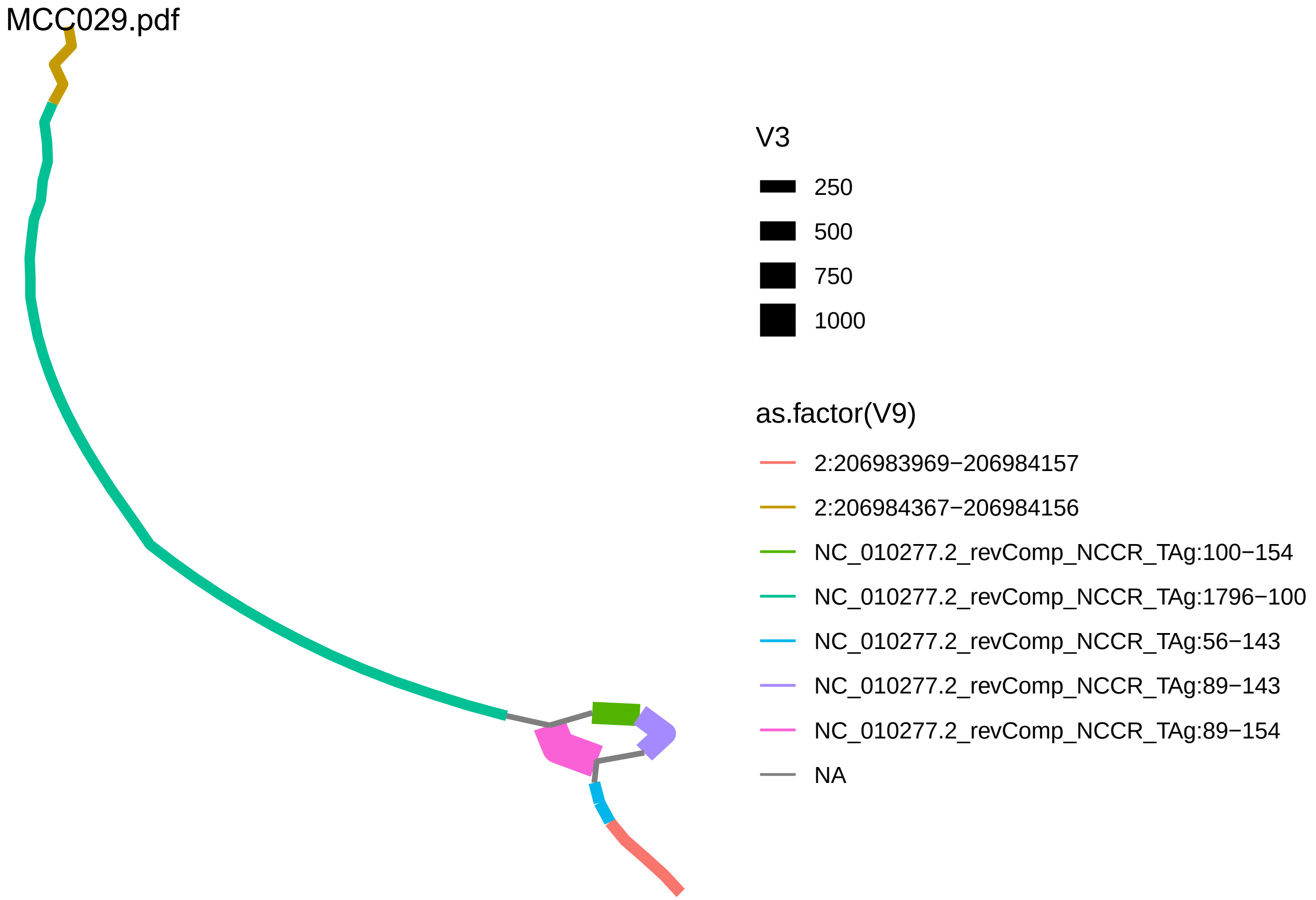
MCC029.pdf
V3
250
500
750
1000
as.factor(V9)
2:206983969−206984157
2:206984367−206984156
NC_010277.2_revComp_NCCR_TAg:100−154
NC_010277.2_revComp_NCCR_TAg:1796−100
NC_010277.2_revComp_NCCR_TAg:56−143
NC_010277.2_revComp_NCCR_TAg:89−143
NC_010277.2_revComp_NCCR_TAg:89−154
NA

MCC036.pdf

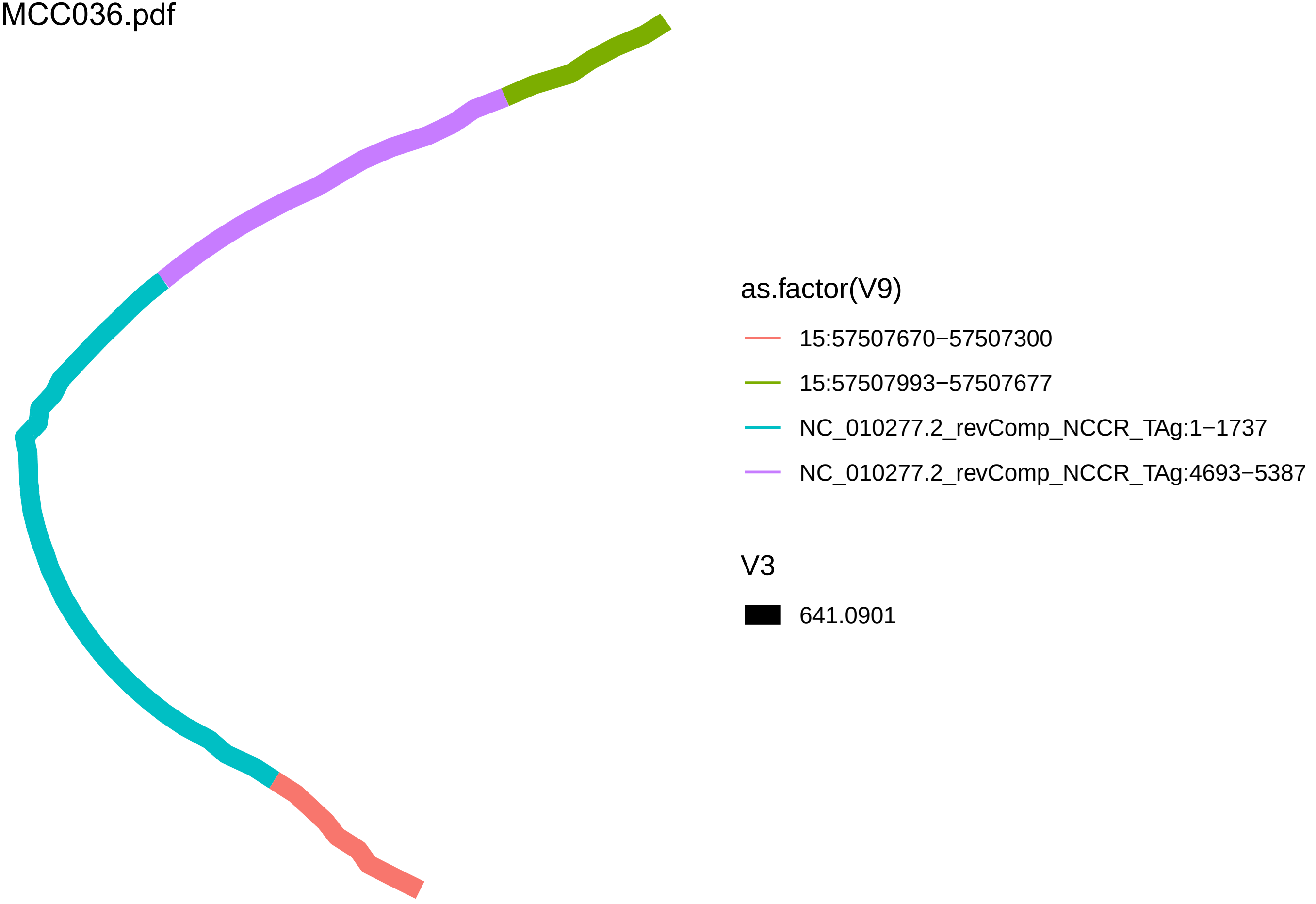

as.factor(V9)
15:57507670−57507300
15:57507993−57507677
NC_010277.2_revComp_NCCR_TAg:1−1737
NC_010277.2_revComp_NCCR_TAg:4693−5387
V3
641.0901

MCC037.pdf

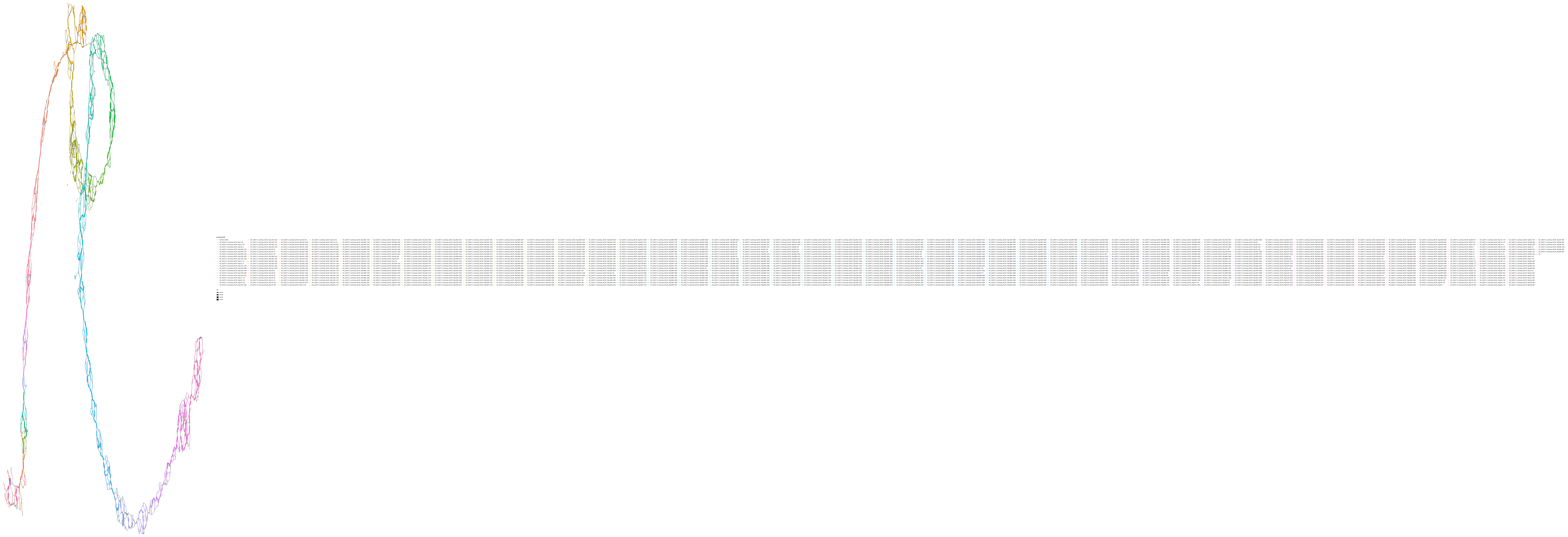

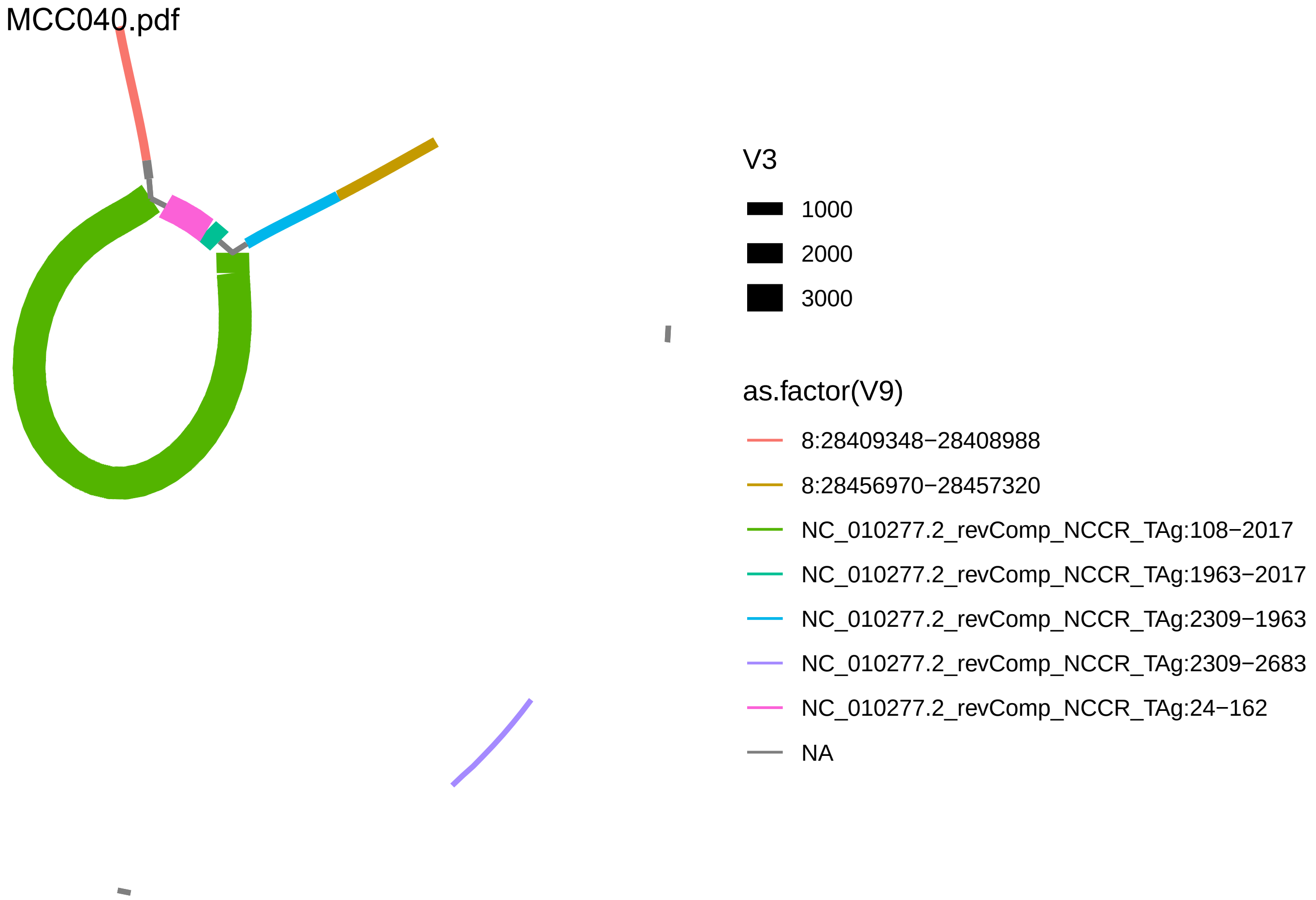
MCC040.pdf
V3
1000
2000
3000
as.factor(V9)
8:28409348−28408988
8:28456970−28457320
NC_010277.2_revComp_NCCR_TAg:108−2017
NC_010277.2_revComp_NCCR_TAg:1963−2017
NC_010277.2_revComp_NCCR_TAg:2309−1963
NC_010277.2_revComp_NCCR_TAg:2309−2683
NC_010277.2_revComp_NCCR_TAg:24−162
NA

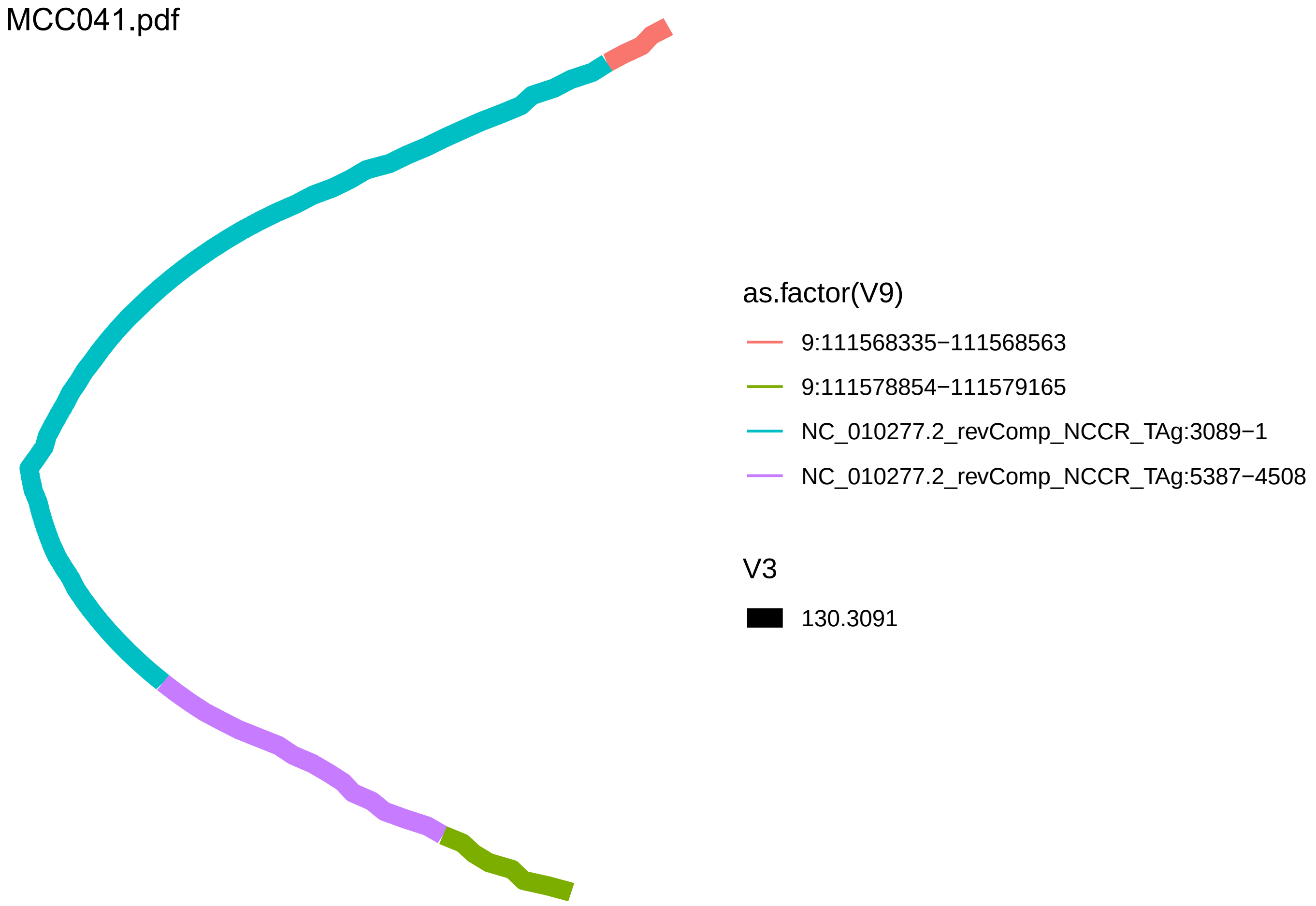
MCC041.pdf
as.factor(V9)
9:111568335−111568563
9:111578854−111579165
NC_010277.2_revComp_NCCR_TAg:3089−1
NC_010277.2_revComp_NCCR_TAg:5387−4508
V3
130.3091

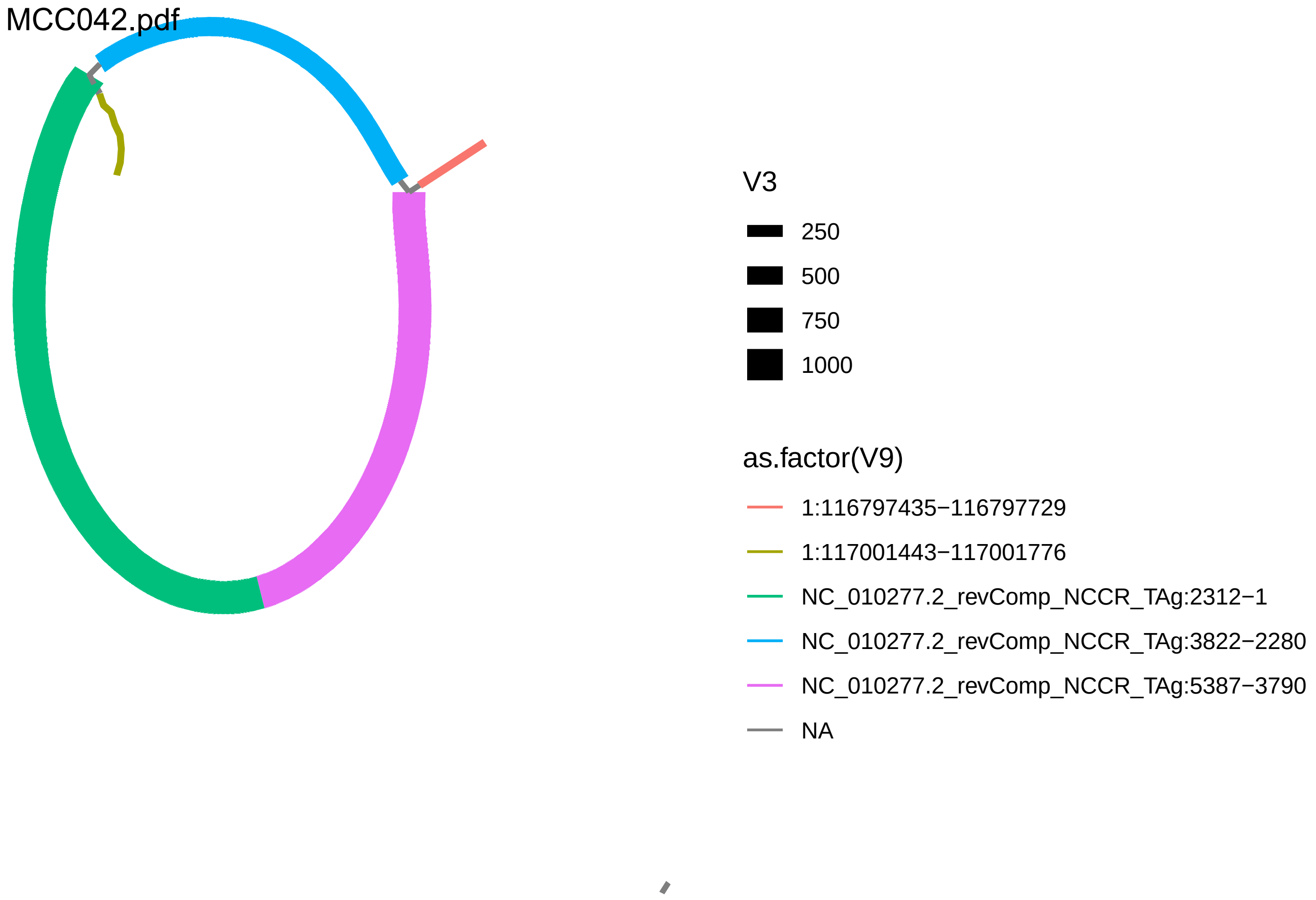

MCC042.pdf
V3
250
500
750
1000
as.factor(V9)
1:116797435−116797729
1:117001443−117001776
NC_010277.2_revComp_NCCR_TAg:2312−1
NC_010277.2_revComp_NCCR_TAg:3822−2280
NC_010277.2_revComp_NCCR_TAg:5387−3790
NA

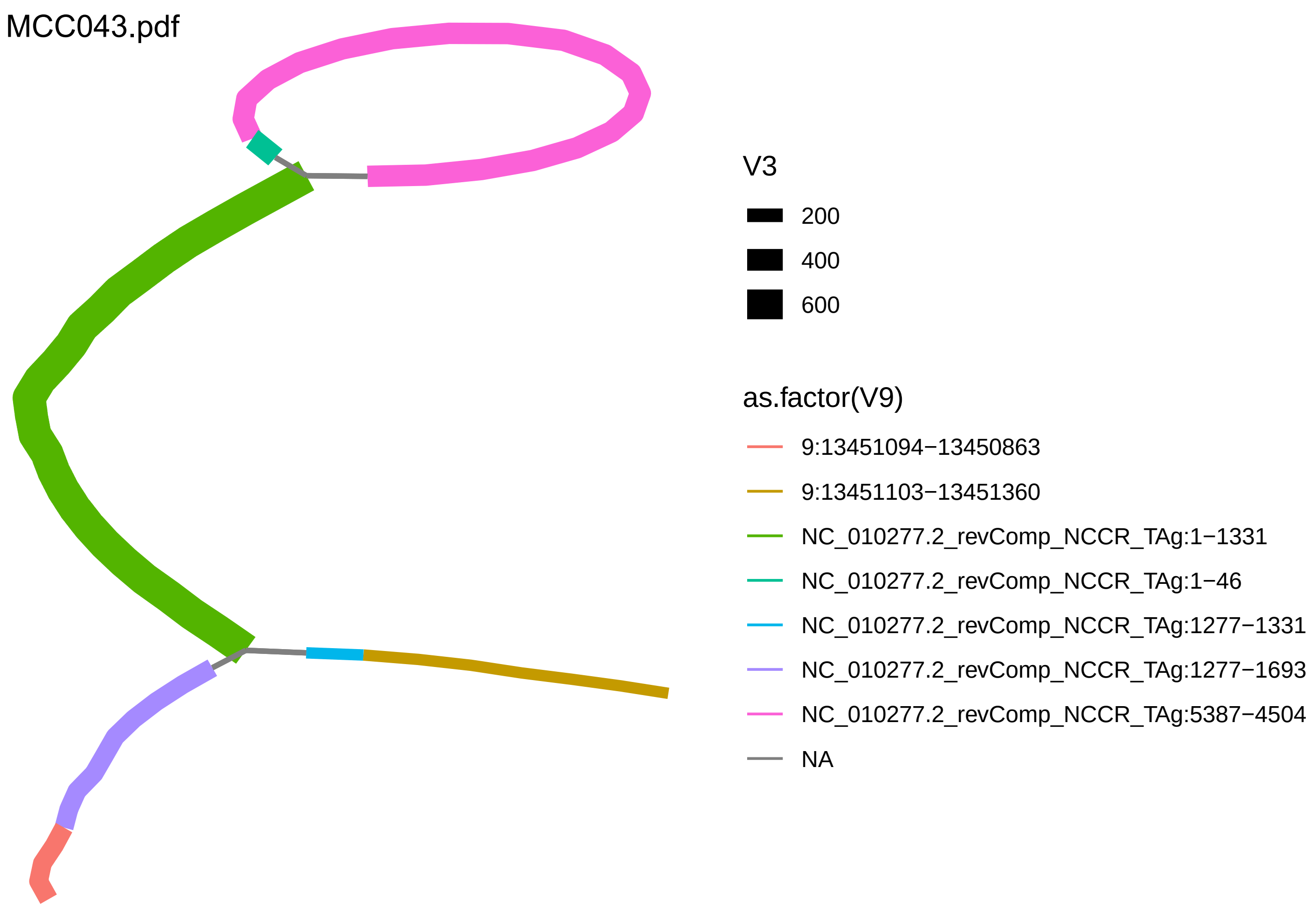
MCC043.pdf
V3
200
400
600
as.factor(V9)
9:13451094−13450863
9:13451103−13451360
NC_010277.2_revComp_NCCR_TAg:1−1331
NC_010277.2_revComp_NCCR_TAg:1−46
NC_010277.2_revComp_NCCR_TAg:1277−1331
NC_010277.2_revComp_NCCR_TAg:1277−1693
NC_010277.2_revComp_NCCR_TAg:5387−4504
NA

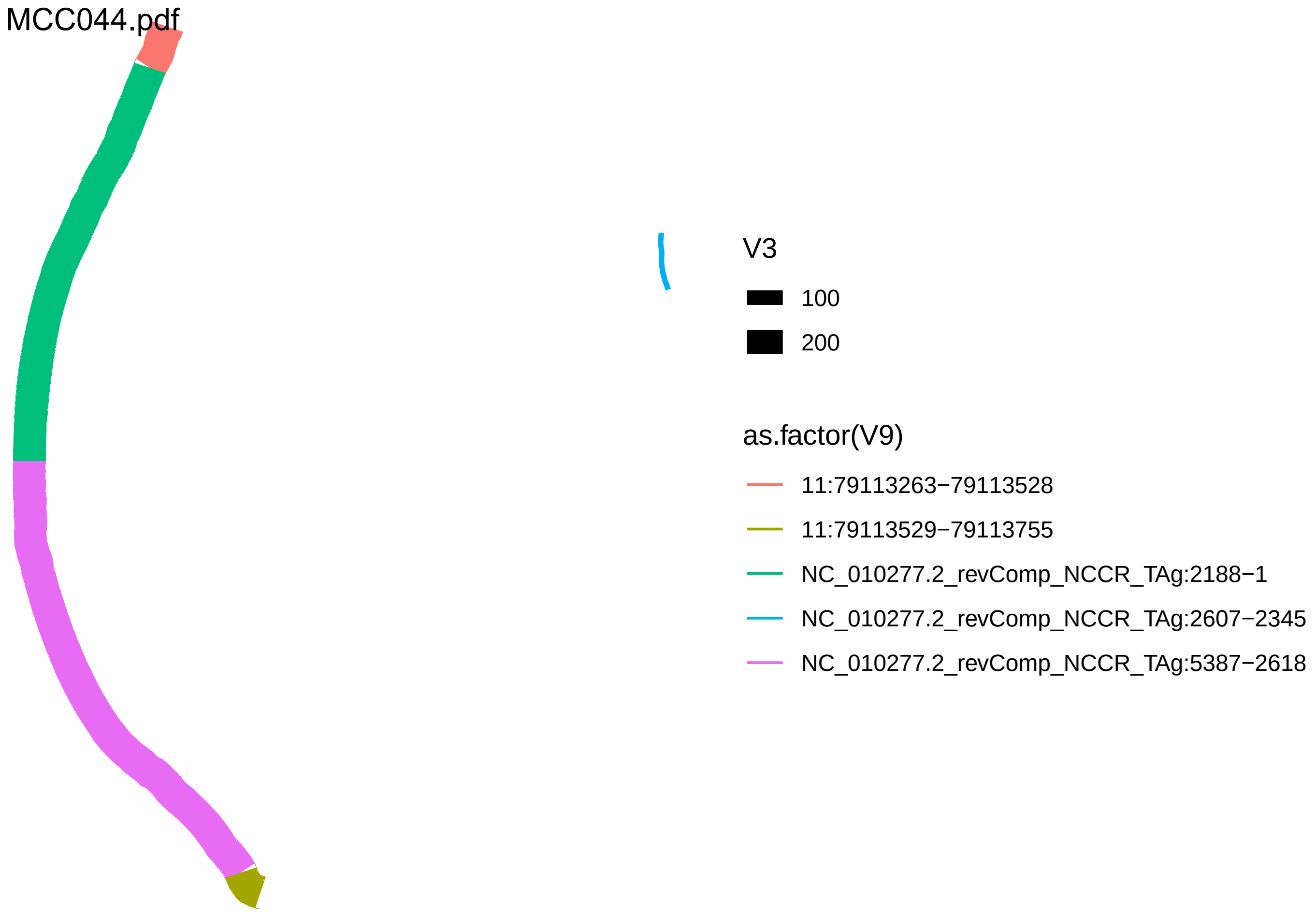
MCC044.pdf
V3
100
200
as.factor(V9)
11:79113263−79113528
11:79113529−79113755
NC_010277.2_revComp_NCCR_TAg:2188−1
NC_010277.2_revComp_NCCR_TAg:2607−2345
NC_010277.2_revComp_NCCR_TAg:5387−2618

MCC047.pdf

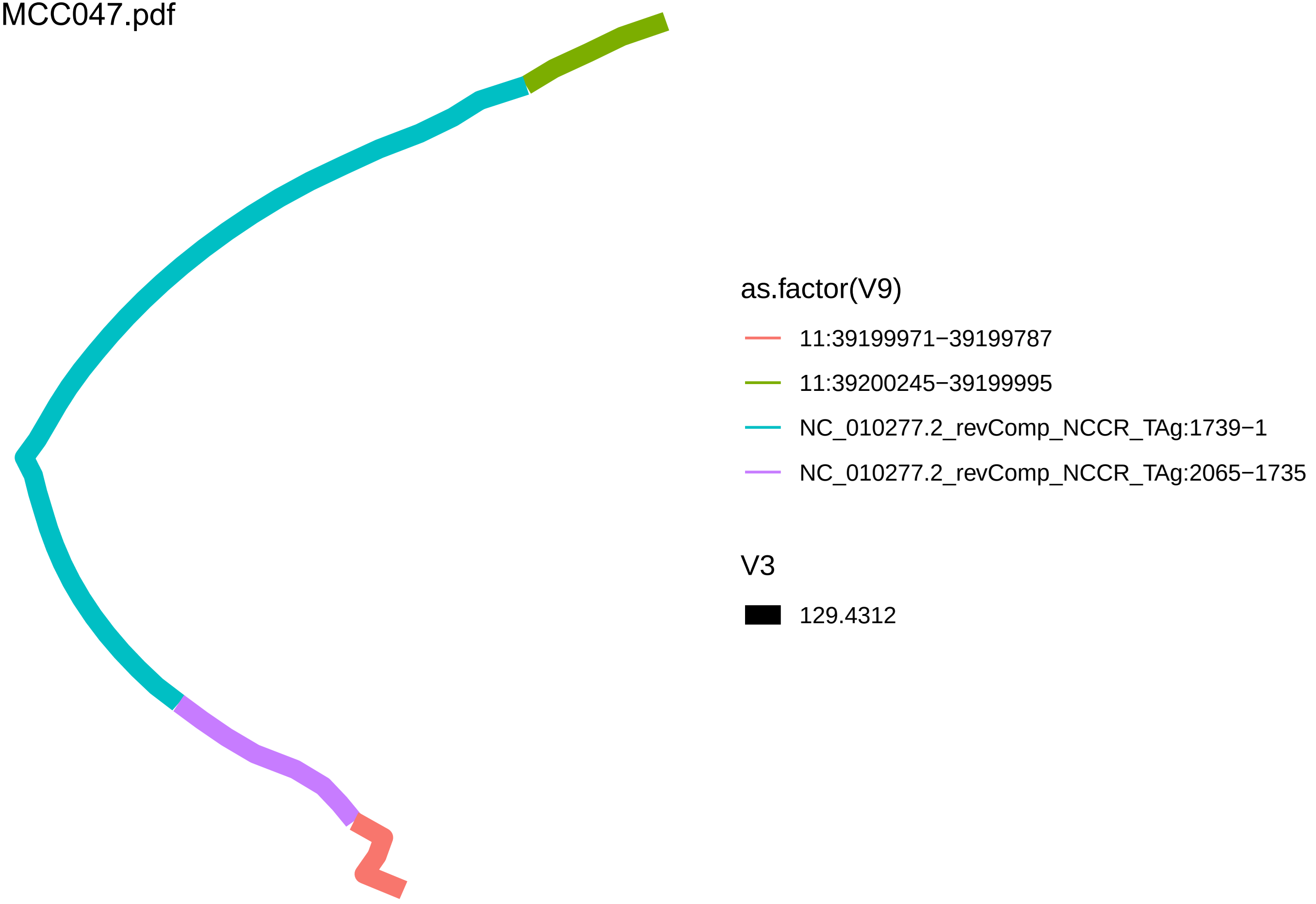
as.factor(V9)
11:39199971−39199787
11:39200245−39199995
NC_010277.2_revComp_NCCR_TAg:1739−1
NC_010277.2_revComp_NCCR_TAg:2065−1735
V3
129.4312

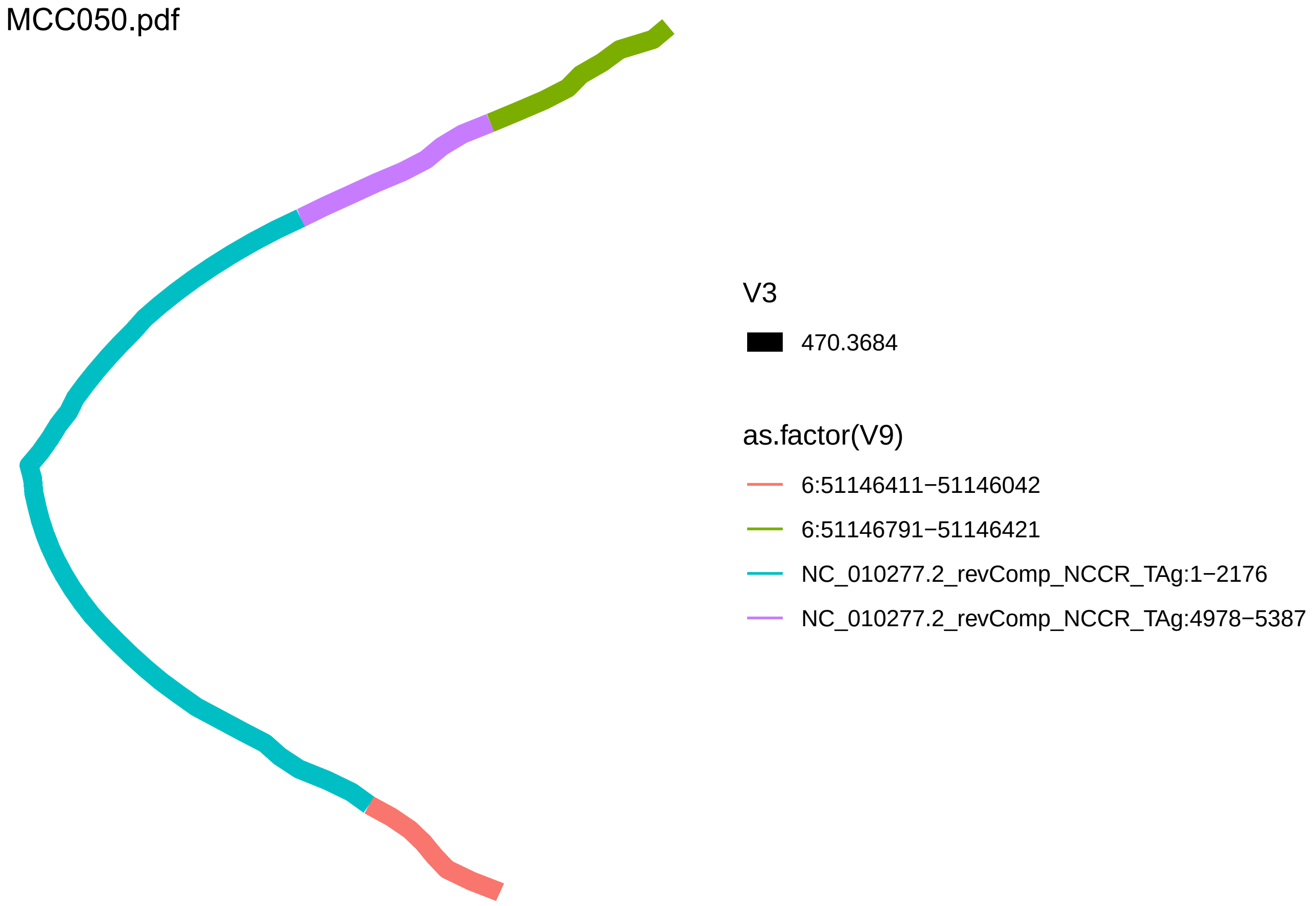

MCC050.pdf
V3
470.3684
as.factor(V9)
6:51146411−51146042
6:51146791−51146421
NC_010277.2_revComp_NCCR_TAg:1−2176
NC_010277.2_revComp_NCCR_TAg:4978−5387

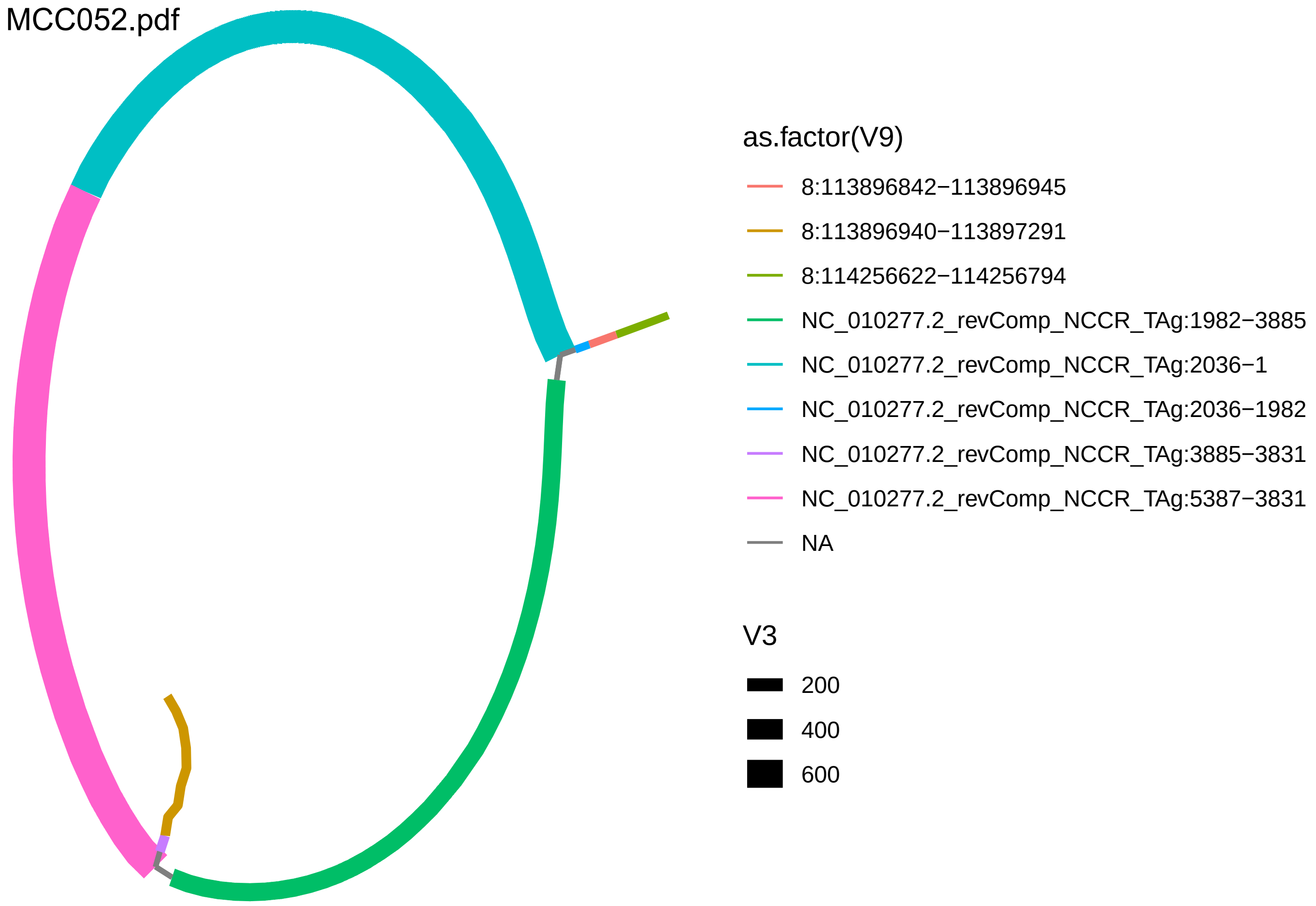
MCC052.pdf
as.factor(V9)
8:113896842−113896945
8:113896940−113897291
8:114256622−114256794
NC_010277.2_revComp_NCCR_TAg:1982−3885
NC_010277.2_revComp_NCCR_TAg:2036−1
NC_010277.2_revComp_NCCR_TAg:2036−1982
NC_010277.2_revComp_NCCR_TAg:3885−3831
NC_010277.2_revComp_NCCR_TAg:5387−3831
NA
V3
200
400
600

MCC054.pdf

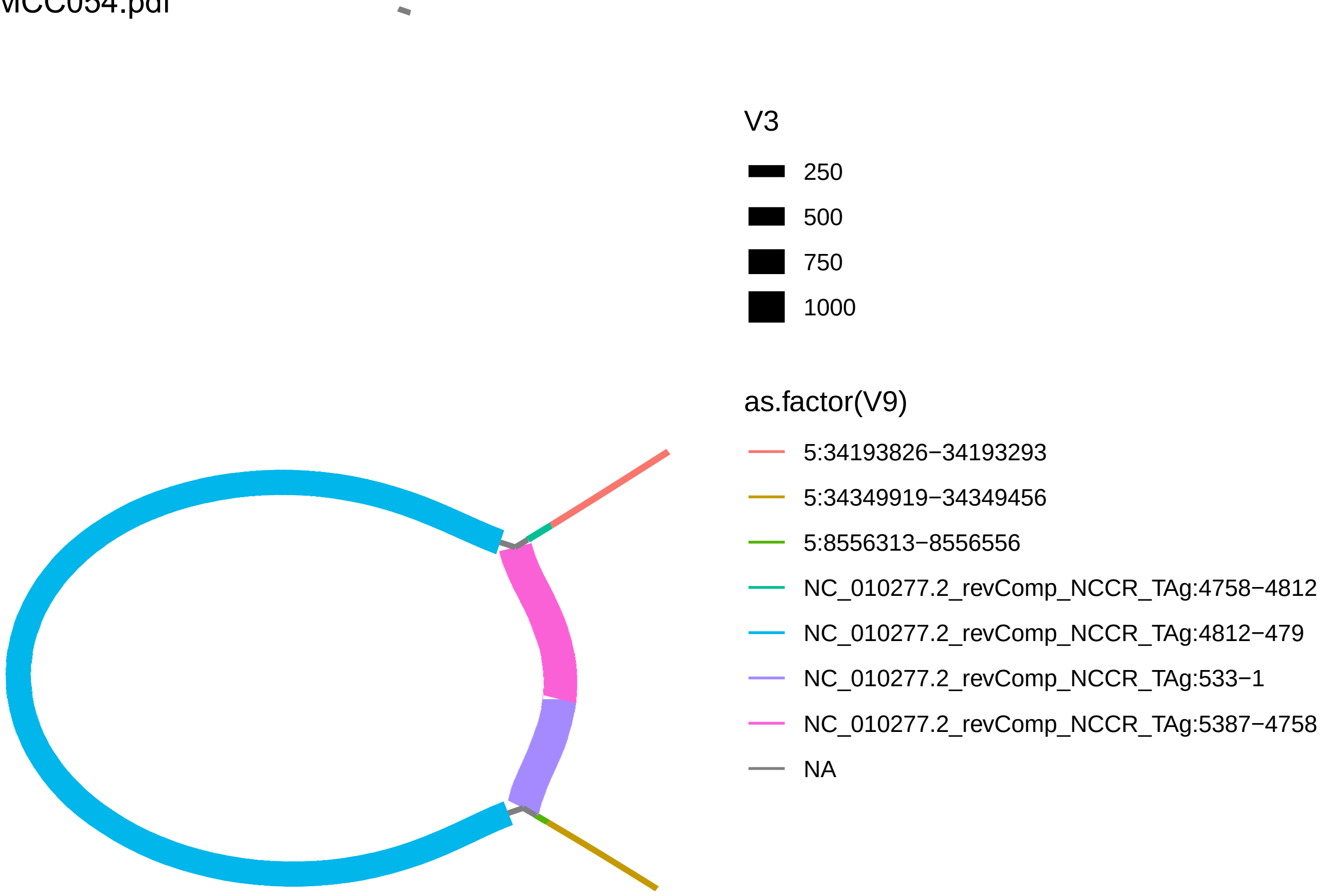
V3
250
500
750
1000
as.factor(V9)
5:34193826−34193293
5:34349919−34349456
5:8556313−8556556
NC_010277.2_revComp_NCCR_TAg:4758−4812
NC_010277.2_revComp_NCCR_TAg:4812−479
NC_010277.2_revComp_NCCR_TAg:533−1
NC_010277.2_revComp_NCCR_TAg:5387−4758
NA

MCC056.pdf

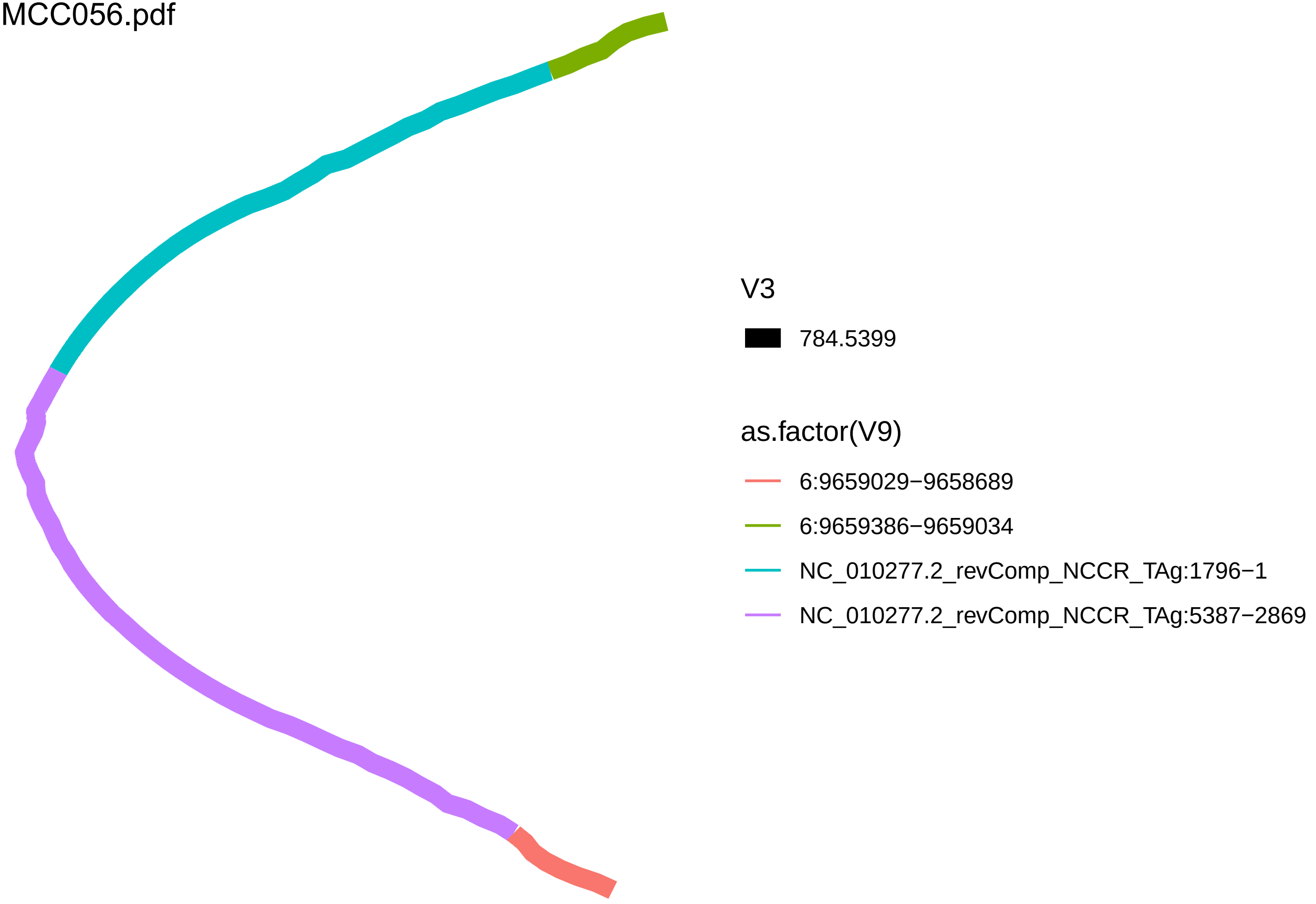
V3
784.5399
as.factor(V9)
6:9659029−9658689
6:9659386−9659034
NC_010277.2_revComp_NCCR_TAg:1796−1
NC_010277.2_revComp_NCCR_TAg:5387−2869

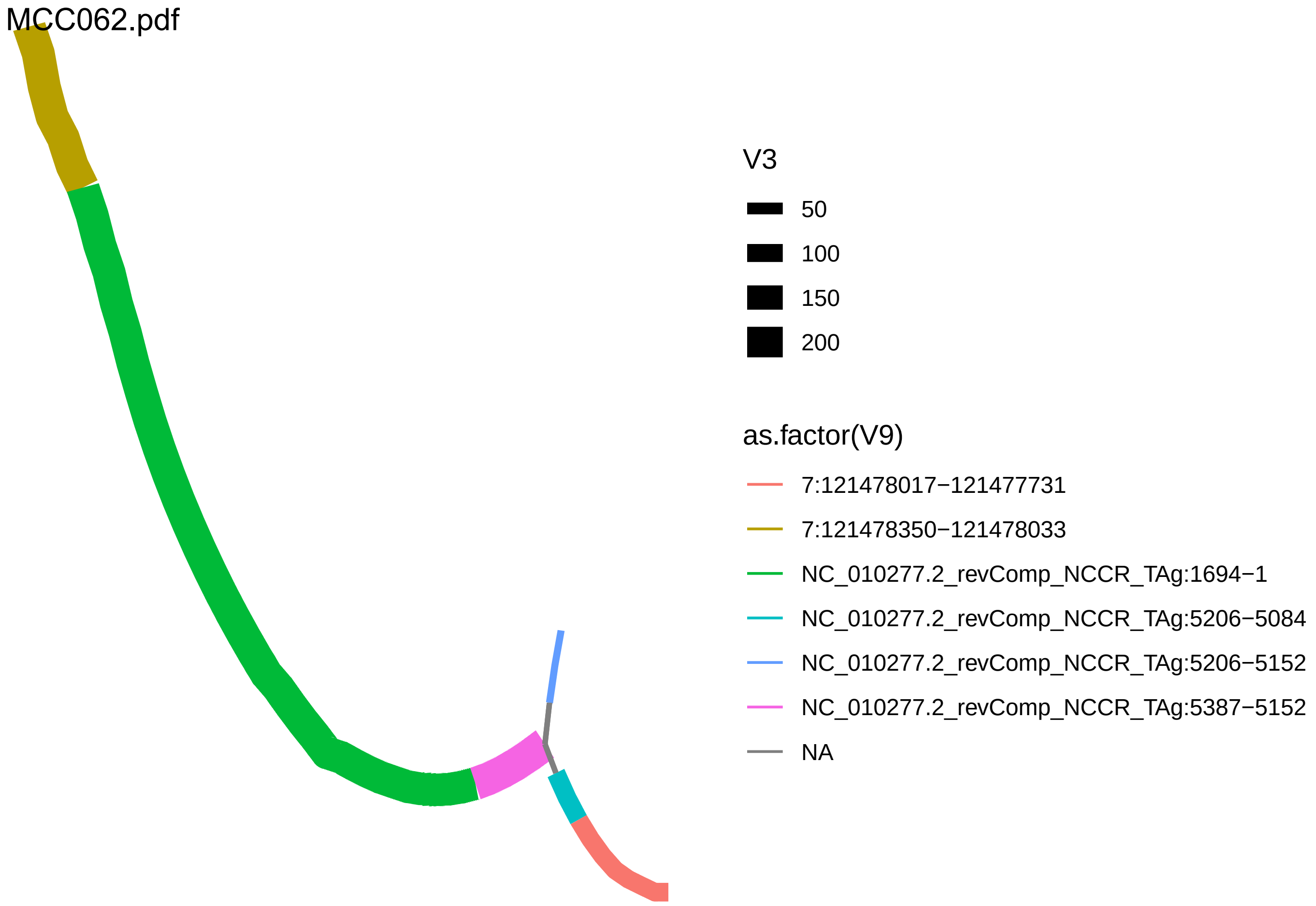
MCC062.pdf
V3
50
100
150
200
as.factor(V9)
7:121478017−121477731
7:121478350−121478033
NC_010277.2_revComp_NCCR_TAg:1694−1
NC_010277.2_revComp_NCCR_TAg:5206−5084
NC_010277.2_revComp_NCCR_TAg:5206−5152
NC_010277.2_revComp_NCCR_TAg:5387−5152
NA

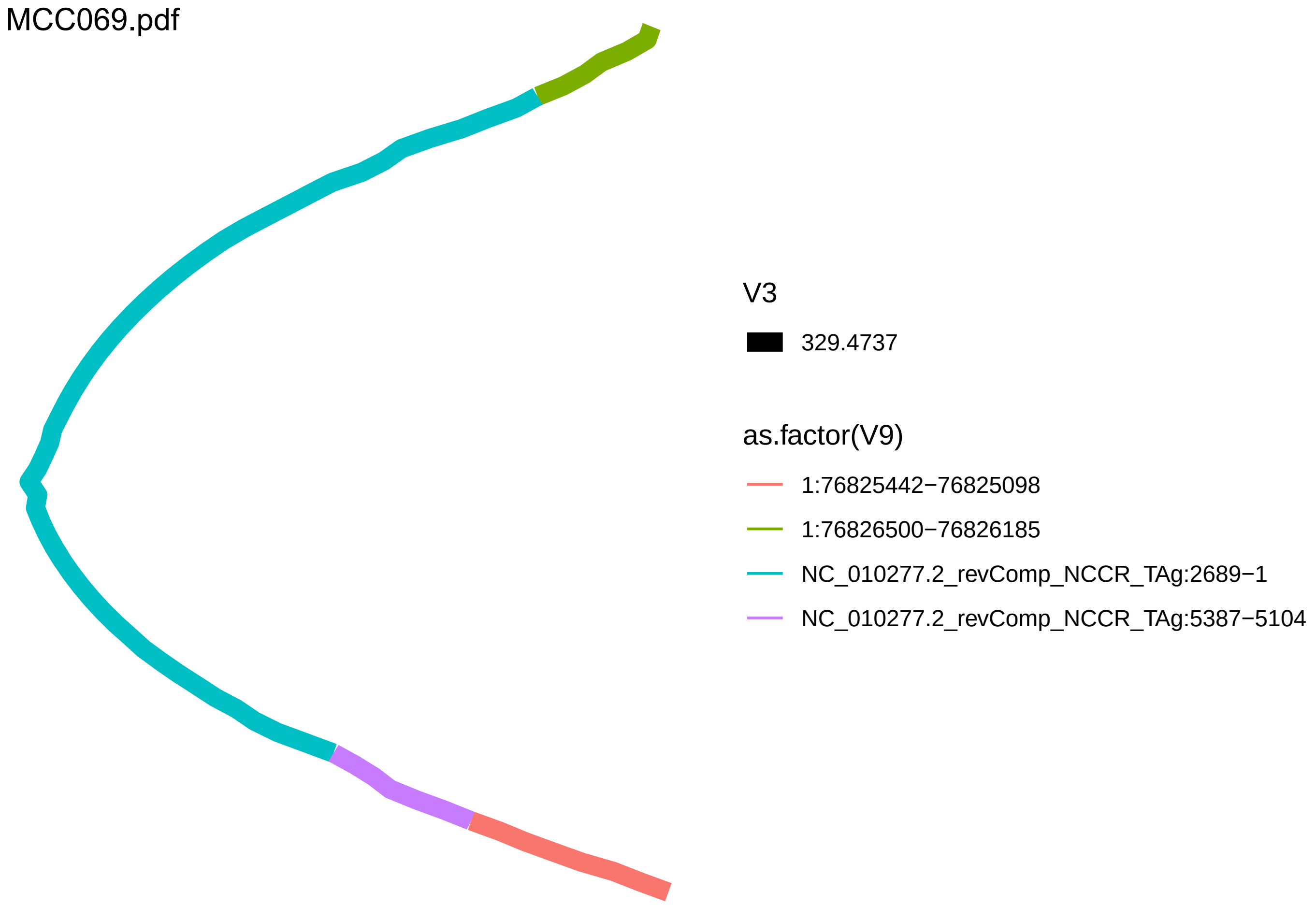
MCC069.pdf
V3
329.4737
as.factor(V9)
1:76825442−76825098
1:76826500−76826185
NC_010277.2_revComp_NCCR_TAg:2689−1
NC_010277.2_revComp_NCCR_TAg:5387−5104

MCC071.pdf

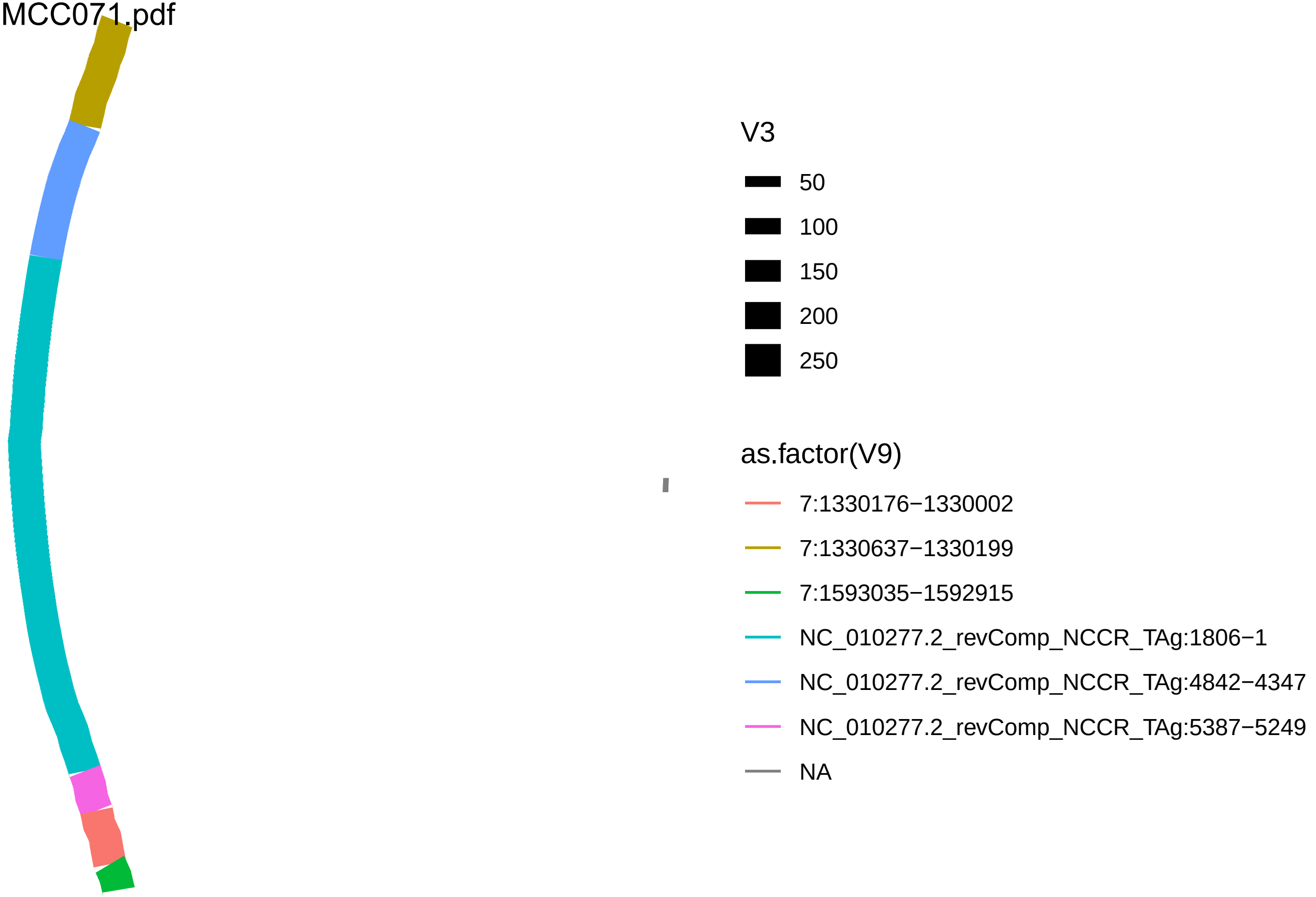
V3
50
100
150
200
250
as.factor(V9)
7:1330176−1330002
7:1330637−1330199
7:1593035−1592915
NC_010277.2_revComp_NCCR_TAg:1806−1
NC_010277.2_revComp_NCCR_TAg:4842−4347
NC_010277.2_revComp_NCCR_TAg:5387−5249
NA
